# The within-subject application of diffusion tensor MRI and CLARITY reveals brain structural changes in *Nrxn2* deletion mice

**DOI:** 10.1101/300806

**Authors:** Eleftheria Pervolaraki, Adam L. Tyson, Francesca Pibiri, Steven L. Poulter, Amy C. Reichelt, R. John Rodgers, Steven J. Clapcote, Colin Lever, Laura C. Andreae, James Dachtler

**Affiliations:** School of Biomedical Sciences, University of Leeds, LS2 9JT, UK; Centre for Developmental Neurobiology, Institute of Psychiatry, Psychology and Neuroscience, King’s College London, London SE1 1UL, UK; MRC Centre for Neurodevelopmental Disorders, King’s College London, London SE1 1UL, UK; Department of Forensic and Neurodevelopmental Sciences, Institute of Psychiatry, Psychology and Neuroscience, King’s College London, London SE5 8AF, UK; Department of Psychology, Durham University, South Road, Durham, DH1 3LE, UK; Robarts Research Institute, Western University, London, Ontario, Canada, N6A 5B7, Canada; School of Psychology, University of Leeds, LS2 9JT, UK

**Keywords:** MRI, CLARITY, social, autism, axons, diffusion, structure, imaging

## Abstract

**Background:** Of the many genetic mutations known to increase the risk of autism spectrum disorder, a large proportion cluster upon synaptic proteins. One such family of presynaptic proteins are the neurexins (NRXN), and recent genetic and mouse evidence has suggested a causative role for *NRXN2* in generating altered social behaviours. Autism has been conceptualised as a disorder of atypical connectivity, yet how single-gene mutations affect such connectivity remains under-explored. To attempt to address this, we have developed a quantitative analysis of microstructure and structural connectivity leveraging diffusion tensor MRI (DTI) with high-resolution 3D imaging in optically cleared (CLARITY) brain tissue in the same mouse, applied here to the *Nrxn2α* knockout (KO) model.

**Methods:** Fixed brains of *Nrxn2α* KO mice underwent DTI using 9.4T MRI, and diffusion properties of socially-relevant brain regions were quantified. The same tissue was then subjected to CLARITY to immunolabel axons and cell bodies, which were also quantified.

**Results:** DTI revealed increases in fractional anisotropy in the amygdala (including the basolateral nuclei), the anterior cingulate cortex, the orbitofrontal cortex and the hippocampus. Axial diffusivity of the anterior cingulate cortex and orbitofrontal cortex was significantly increased in *Nrxn2α* KO mice, as were tracts between the amygdala and the orbitofrontal cortex. Using CLARITY, we find significantly altered axonal orientation in the amygdala, orbitofrontal cortex and the anterior cingulate cortex, which was unrelated to cell density.

**Conclusions:** Our findings demonstrate that deleting a single neurexin gene (Nrxn2α) induces atypical structural connectivity within socially-relevant brain regions. More generally, our combined within-subject DTI and CLARITY approach presents a new, more sensitive method of revealing hitherto undetectable differences in the autistic brain.

## Background

Autism is a common neurodevelopmental disorder, which is highly heritable (1). While heritability is high, it is also clear that autism is highly polygenic. Around ∼400-1000 genes are involved in autism susceptibility (2-5). Many of these genes cluster upon proteins relating to synaptic signaling (6). A family of presynaptic proteins garnering recent interest have been the neurexins (*NRXNs*). NRXNs are encoded by three genes (*NRXN1, NRXN2, NRXN3;* note that *CNTNAP1* and *CNTNAP2* are sometimes referred to as *NRXN4*), of which two major isoforms exist: the longer α proteins with six laminin/neurexin/sex hormone (LNS) binding domains, and the shorter β proteins with one LNS binding domain (7, 8).

Mutations within all three *NRXN* genes have been linked to autism (6). Heterozygous deletions within *NRXN2* have been identified in a number of individuals with autistic phenotypes. These include an autistic boy and his father (who had severe language delay but not autism) who both had a frameshift mutation within exon 12 of *NRXN2* (9); a 570-kb de novo deletion of 24 genes at chromosome 11q13.1, including *NRXN2*, in a 21-year old man displaying a clinical phenotype including autistic traits (10); a 1.6Mb deletion at chromosome region 11q12.3-11q13.1, including *NRXN2*, in a 23-year old man with intellectual disability and behavioral problems (11); a de novo frameshift mutation identified in a Chinese man with autism spectrum disorder (ASD) (12), a 921 kb microdeletion at 11q13 in a 2 year old boy who had language and developmental delay (although did not meet the autism diagnosis criteria) (13) and a paternally inherited microRNA miR-873-5p variant in an ASD individual which altered binding affinity for several risk-genes including *NRXN2* and *CNTNAP2* (*NRXN4*) (14). Furthermore, recently, two large-scale reports have identified *NRXN2* with ASD risk. A study of 529 ASD patients and 1,923 controls in a Chinese population identified two *NRXN2* variants which significantly increase ASD risk (15). The second study employed machine learning approaches across 5000 ASD families to rank the importance of ASD candidate genes, and ranks *NRXN2* in the top ∼0.5% of genes, i.e. 113^th^ (16). For comparison, *NRXN1*, for which the evidence base for its links to ASD is broader and stronger, ranks 45, and *CNTNAP2* ranks 211^th^ (16). Consistent with these association studies, we and others have previously found that homozygous or heterozygous deletion of *Nrxn2α* induces impairment in social approach and social recognition (17-19). In summary, although mutations within *NRXN2* are rare, understanding how they may drive social, ASD-relevant behavioural changes is important. One important goal is to help elucidate how apparently convergent pathophysiology in ASD emerges despite marked genetic heterogeneity (Insert ref Geschwind & State, 2015 cited above); mapping brain alterations driven by different single genes is thus a crucial task.

Currently it is unknown whether deletion of *Nrxn2α* changes the brain’s microstructure and connectivity. One previous study found coarse alterations to cell layer thickness within the hippocampus of *Nrxn2α* homozygous KOs (20). However, cell density measurements are unlikely to reveal the true extent of changes within the autistic brain. Within the current study, we have addressed this by developing a dual imaging approach (DTI and CLARITY) that quantifies the alignment and density of white matter, applied here to brain regions known to support social behavior in a mouse model of autism.

Diffusion tensor MRI (or DTI) is based upon the movement of water molecules, a measure that is termed fractional anisotropy (FA). Apparent diffusion coefficient (ADC) is similar to FA, but quantifies diffusion restriction as opposed to the spatial symmetry of diffusion. This approach has been used to explore neuropathological markers in autistic patients; alterations in myelination, axonal abundance, size and orientation all modify FA and ADC values (21-23). Using the preferred direction of the diffusion of tensors between brain regions can be used to explore their potential connection.

Quantification of those computed streamlines by FA and axial and/or radial diffusion can indicate impairments in regional structural connectivity. Since aberrant brain connectivity is likely a core feature of autism (24), we reasoned that the candidate method for probing the autistic brain should combine tractographic techniques. Accordingly, here, we combined high resolution imaging of labelled neuronal tracts in brains rendered transparent by CLARITY with DTI.

CLARITY is a recent development that renders tissue optically transparent and macromolecule permeable (25). This permits antibody staining and imaging of much larger tissue volumes than possible under traditional immunofluorescence techniques. By examining fiber orientation without sectioning-related artefacts and biases, axonal staining in cleared tissue affords a deeper understanding of the microstructure and structural connectivity of a brain region.

Given the social impairments found within *Nrxn2α* mice, we sought to examine those brain regions most closely linked with social behavior (See Supplemental Materials). Briefly, we identified four regions of interest (ROIs): the amygdala, and three brain regions strongly and directly connected to the amygdala; the hippocampus, orbitofrontal cortex (OFC), and anterior cingulate cortex (ACC). As predicted, structural connectivity was abnormal in *Nrxn2α* mice.

## Methods

### Ethics

All procedures were approved by the University of Leeds and Durham University Animal Ethical and Welfare Review Boards and were performed under UK Home Office Project and Personal Licenses in accordance with the Animals (Scientific Procedures) Act 1986.

### Animals

Full details of the animals, their background, genotyping and housing can be found elsewhere (17). In brief, male B6;129-*Nrxn3tm1Sud*/*Nrxn1tm1Sud*/*Nrxn2tm1Sud*/J mice (JAX #006377) were purchased from the Jackson Laboratory and outbred once to the C57BL/6NCrl strain (Charles River, Margate, United Kingdom) to obtain mice that were individually *Nrxn2α* KO heterozygotes. Subsequently, HET knockout males were bred with HET females (cousin mating).

### Experimental animals

6 adult wild-type males (Charles River, Margate, UK) and 6 age matched littermate *Nrxn2α* KO homozygotes (71 days ± 6 days old (SEM)) were perfused-fixed with 4% paraformaldehyde (PFA) in 0.1 M phosphate buffer saline (PBS) and the brains extracted. The brains were immersed in 4% PFA/0.1 M PBS for a minimum of 48 hours prior to imaging. Mouse weights were no specifically taken prior to perfusion. However, in a separate cohort, wild-type and *Nrxn2α* KO homozygotes did not significantly differ in body mass (wild-type, n = 15, 30.9 ± 4.1 g; *Nrxn2* KO, n = 10, 28.6 ± 4.3 g, t-test p = 0.167). We did not specifically time perfusions, although as a matter of process, each mouse was perfused with ∼60 ml of fixative. We cannot rule out that variance in perfusion timings may have influenced the results, which is a limitation of the current study. During imaging, the samples were placed in custom-built MR-compatible tubes containing Fomblin Y (Sigma, Poole, Dorset, UK).

Due to the relatively low variance, and owing to the complexity and methodological nature in our experimental approach, we achieved significance by groups of 6 (power provided in Results). No data was excluded from the study. Sample randomisation was performed by JD, with experimenters (EP and ALT) blinded to genotype.

### Data Acquisition

Image acquisition has been described elsewhere (26). Each brain was 3D imaged using the protocol TE: 35 ms, TR: 700 ms and 10 signal averages. The field of view was set at 128 × 128 × 128, with a cubic resolution of 100 μm/pixel and a b value of 1200 s/mm^2^. Further details can be found in Supplemental Materials.

### Image Processing

Parsing of the raw data was semi-automated using DSI Studio, in order to obtain b-values for every normalized gradient vector on the x, y and z orientations. Unwanted background, setting a threshold, smoothing of the data and definition of tissue boundaries was performed prior to the reconstruction of the final 3D image. DTI analysis parameters were calculated as previously described (27).

The *ex vivo* mouse brain 3D diffusion-weighted images were reconstructed from the Bruker binary file using DSI Studio (http://dsi-studio.labsolver.org) (28). Direction Encoded Colour Map (DEC) images were generated by combining the information from the primary eigenvectors, diffusion images and the FA. Images of the primary vectors and their orientation were reconstructed and superimposed on corresponding FA images to guide the segmentation of discrete anatomical locations according to the brain atlas (Figure 1B-D). Region of interest definition was performed by author EP and corroborated independently by JD, with region area compared between the experimenters (data not shown). For whole brain region analysis, we used a similar approach, except regions were segmented for every other slice in the anterior to posterior extent (Figure 1A-D; Supp. Figure 1) (29). The DSI Studio DTI reconstruction characterizes the major diffusion direction of the fibre within the brain (30, 31). Extraction of FA (calculated (26)) and ADC was performed within selected segmented brain areas for every 3D reconstructed mouse brain.

**Figure 1.**
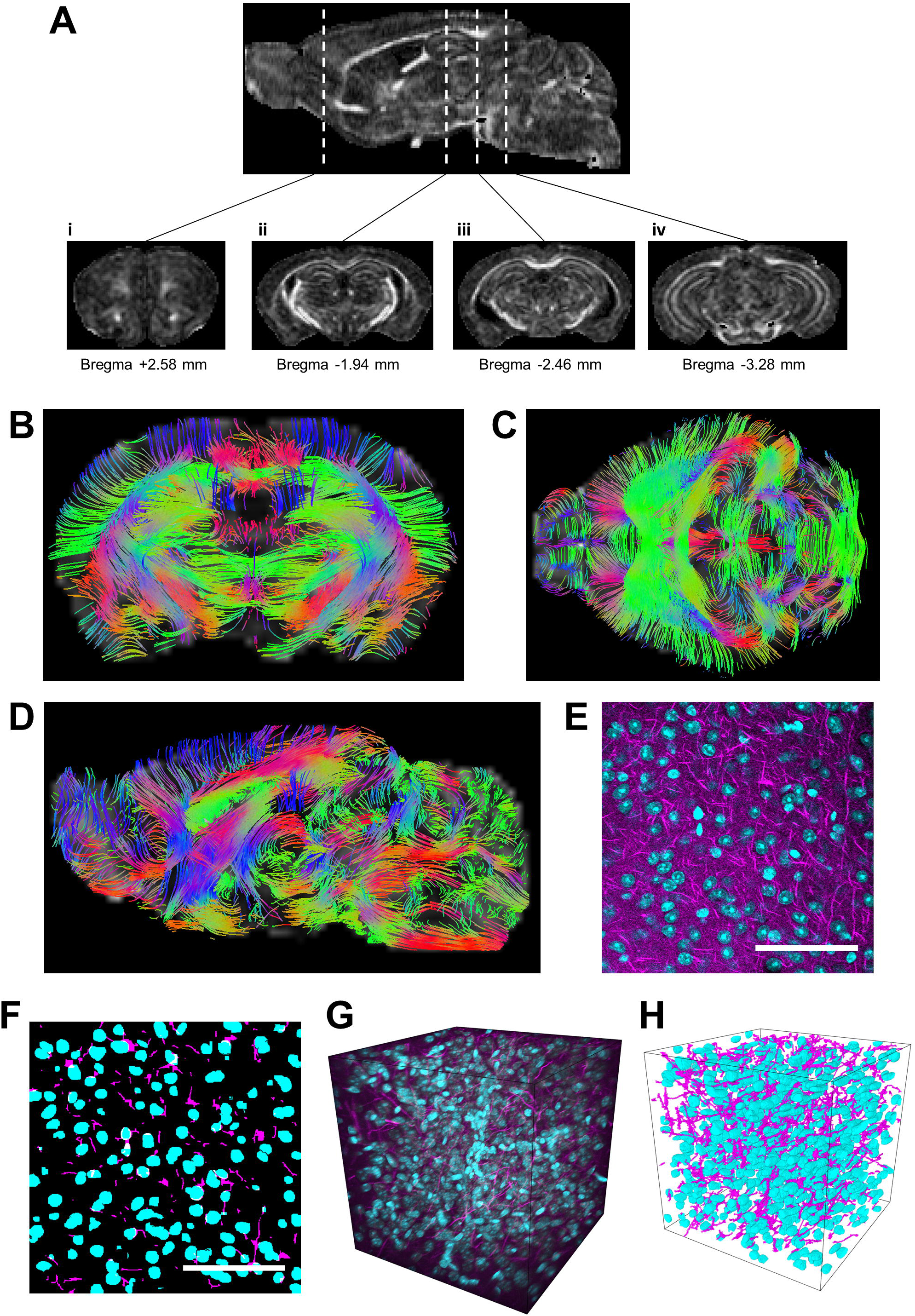
Quantification of CLARITY imaging. **A** Sections of DTI-scanned brain were segmented at different Bregma levels for (i) the orbitofrontal cortex, (ii) the anterior hippocampus and amygdala, (iii) the mid hippocampus and posterior amygdala and (iv) the posterior hippocampus. **B-D** DTI-scanned brains were computed for tracts. Tissue from wild-type and *Nrxn2α* KO mice were cleared and stained for neurofilament and DAPI (**E**). **F** Automated MATLAB scripts were used to segment the DAPI (blue) and neurofilament (purple) channels such that cell density and axonal density and orientation could be calculated. **G** is representative of a CLARITY-derived 3D stacked image of a DAPI and neurofilament of a region of interest, with **H** being the corresponding segmented image. Scale bar: 100 μm.

### Regions of Interest (ROIs)

Our DTI approach was to undertake an *a posteriori* analysis of neural organization in regions of interest (ROIs) identified by previous literature as socially-relevant. Given the social impairments found within *Nrxn2α* mice, for the current study, we identified the brain regions of interest (ROIs) most closely linked with social behavior, using previously published reports of brain region involvement in social behaviour. Quantification of c-Fos immunoreactivity has highlighted the importance of several amygdala nuclei (including the basolateral) following social exposure (32), but also the anterior cingulate cortex (ACC), prefrontal cortex and the hippocampus (33). Lesions to the primate amygdala alter social interactions (34, 35), and amygdala neurons in primates including humans increase firing rates during social scenarios (36-38). Consistent with these animal studies, amygdala damage in humans (39) and amygdala dysfunction in ASD patients (40) impair social responses. Other socially-important brain regions have also been proposed. Notably, several studies have implicated the rodent hippocampus in social behavior, including social memory and sociability (41-43). For instance, intrahippocampal administration of neurolide-2, which interacts with α-neurexin, specifically impairs sociability, but not anxiety and spatial learning in rats (44). These findings are consistent with reports of social deficits in humans with hippocampal damage (45) and hippocampal abnormalities in ASD (46, 47). Finally, several studies link the frontal cortex, particularly the orbitofrontal cortex, which is strongly anatomically connected with the amygdala(48), to social processing (49, 50), consistent with findings of abnormalities in orbitofrontal cortex in ASD (48, 51). Control regions of the primary motor cortex (M1), primary sensory cortex (S1) and the barrel field were chosen for CLARITY (Supp. Figure 7N-O).

### CLARITY

Following MR imaging, the brains were washed in PBS to remove all Fomblin Y and then incubated for 7 days in hydrogel solution at 4°C prior to polymerisation at 37°C for 3.5 hours. The tissue was cut into 1.5 mm coronal sections using a custom 3D-printed brain-slicing matrix based on MRI scans of an adult C57BL/6 mouse brain (52) and incubated in clearing buffer for 24 days at 37°C with shaking. The cleared tissue was then washed in PBSTN_3_ (0.1% TritonX-100 and 1.5 mM sodium azide in PBS) for 24 hours at room temperature and incubated in primary antibody solution (neurofilament (Aves NF-H) 1:100 in PBSTN_3_) at 37°C with shaking for 13 days. Samples were washed, and then incubated in secondary antibody (AlexaFluor 488 goat anti-chicken IgY) as per the primary. Sections were washed again, and incubated in 3.6 μM DAPI (4’,6-diamidino-2-phenylindole) followed by 85% glycerol in PBS for refractive index matching.

Cleared samples were imaged using a Zeiss 7MP multiphoton microscope at 770 nm using a 20x objective lens (W Plan-Apochromat, NA 1.0, WD 1.7 mm). Images (512 × 512 × 512 voxels or 265 × 265 × 265 µm with an isotropic resolution of 520 nm) were acquired in ACC, basolateral (BLA) and basomedial amygdala and OFC) in both hemispheres. DAPI and neurofilament signal was segmented into cell nuclei and axons, and the resulting binary images were used to generate values for cell density, axonal density and axonal alignment.

Full CLARITY methodological details are available within Supplemental Materials.

### Data Availability

Codes to analyse CLARITY datasets are made available by author LCA by email request to either JD or LCA, subject to reference to the current paper. The datasets used and/or analysed during the current study are available from the corresponding author on reasonable request.

### Data Analysis

All data are expressed as mean ± standard error of the mean (SEM). To assess the variance between genotypes within a single brain structure across hemispheres (given the importance of hemispheric differences in ASD (53)), data was analyzed by within subject repeated measures two-way ANOVAs, with Sidak multiple corrections employed on post hoc testing, or unpaired T-tests. To correct for multiple comparisons, we employed the Benjamini-Hochberg Procedure (corrected P values stated). Non-significant statistical results, particularly hemisphere comparisons, can be found in Supplemental Materials. Statistical testing and graphs were made using GraphPad Prism version 6 and SPSS v22.

## Results

### *Nrxn2α* deletion disrupts DTI measures of microstructure in social brain regions

To assess whether *Nrxn2α* deletion alters gross morphology, we quantified whole brain volume using DTI. We found total brain volume for wild-types and *Nrxn2α* KOs was similar (456.0 ± 14.76 vs. 466.2 ± 11.0 mm^3^ (respectively); t_(10)_ = 0.55, p = 0.59). Thus, *Nrxn2α* deletion does not change total brain size.

To quantitatively measure DTI, we examined FA and ADC. FA analyses changes in the linear orientation (i.e. along an axonal tract), whereas ADC (mean diffusivity) averages diffusion in all directions (i.e. the X, Y and Z axis), which is sensitive to changes such as altered alignment. The amygdala is critically important for social behaviours. To assess whether amygdalar alterations might account for social impairments in *Nrxn2α* KO mice, we segmented the whole amygdala structure and the basolateral nuclei along the anterior-posterior axis.

The amygdala showed a significant increase in FA in *Nrxn2α* KO mice (Figure 2A) (genotype (F_(1, 10)_ = 11.15, p = 0.022, power = 85.2%)). There was a FA reduction was observed specifically in the BLA, a region strongly associated with social behaviours (Figure 2B; genotype (F_(1, 10)_ = 6.31, p = 0.049)). ADC was not significantly altered in the whole amygdala or BLA (Figure 2C&D; all genotype: F_(1, 10)_ <1).

**Figure 2.**
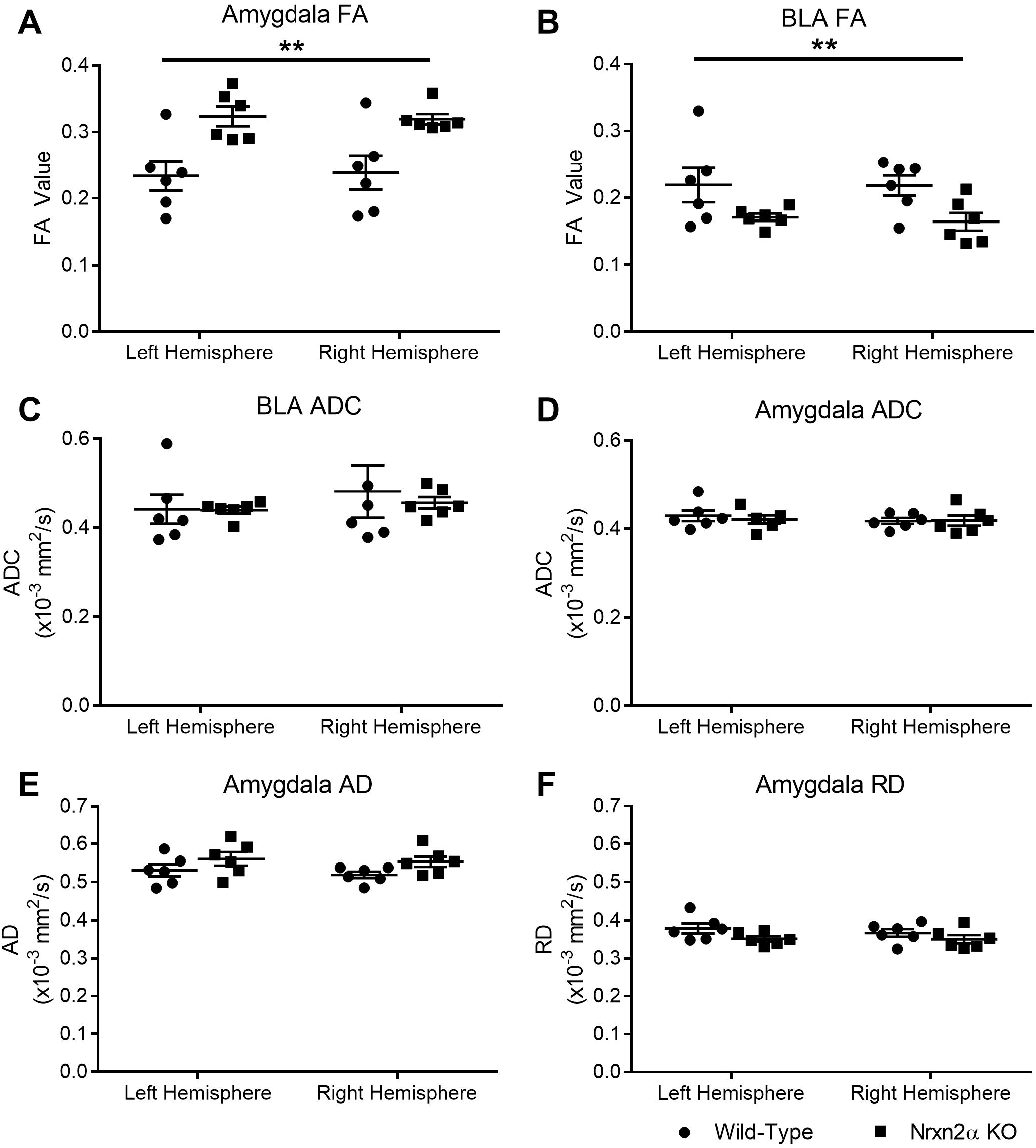
Deletion of *Nrxn2α* increases amygdala fractional anisotropy (FA) but not apparent diffusion coefficient (ADC). DTI images of the amygdala was segmented at two regions; the whole amygdala in the anterior to posterior extent or the basolateral amygdala (BLA) centred at Bregma −1.94 mm. FA of the whole amygdala structure was significantly increased (**A**) but was decreased in the BLA (**B**). However, ADC was similar between the genotypes (**C** and **D**). Axial (AD) (**E**) and radial diffusivity (RD) (**F**) was unaltered in the amygdala. **=P<0.01, *=P<0.05. Error bars represent s.e.m. Wild-type n=6, *Nrxn2α* KO n=6.

We conducted the same analysis for the two prefrontal regions implicated in social behaviour and autism: the OFC and ACC. The pattern of results was similar for both regions: FA was significantly altered, while ADC was unaffected (Figure 3A&B) and the ACC (Figure 3E&F). FA for the OFC was significantly increased (genotype: (F_(1, 10)_ = 16.14, p = 0.009, power = 95.0%)) but ADC was similar between the genotypes (genotype: (F_(1, 10)_ = 1.43, p = 0.11)). The ACC also had significantly increased FA (t_(10)_ = 2.55, p = 0.03, power = 71.0%) but ADC was unaltered (t_(10)_ = 0.51, p = 0.618).

**Figure 3.**
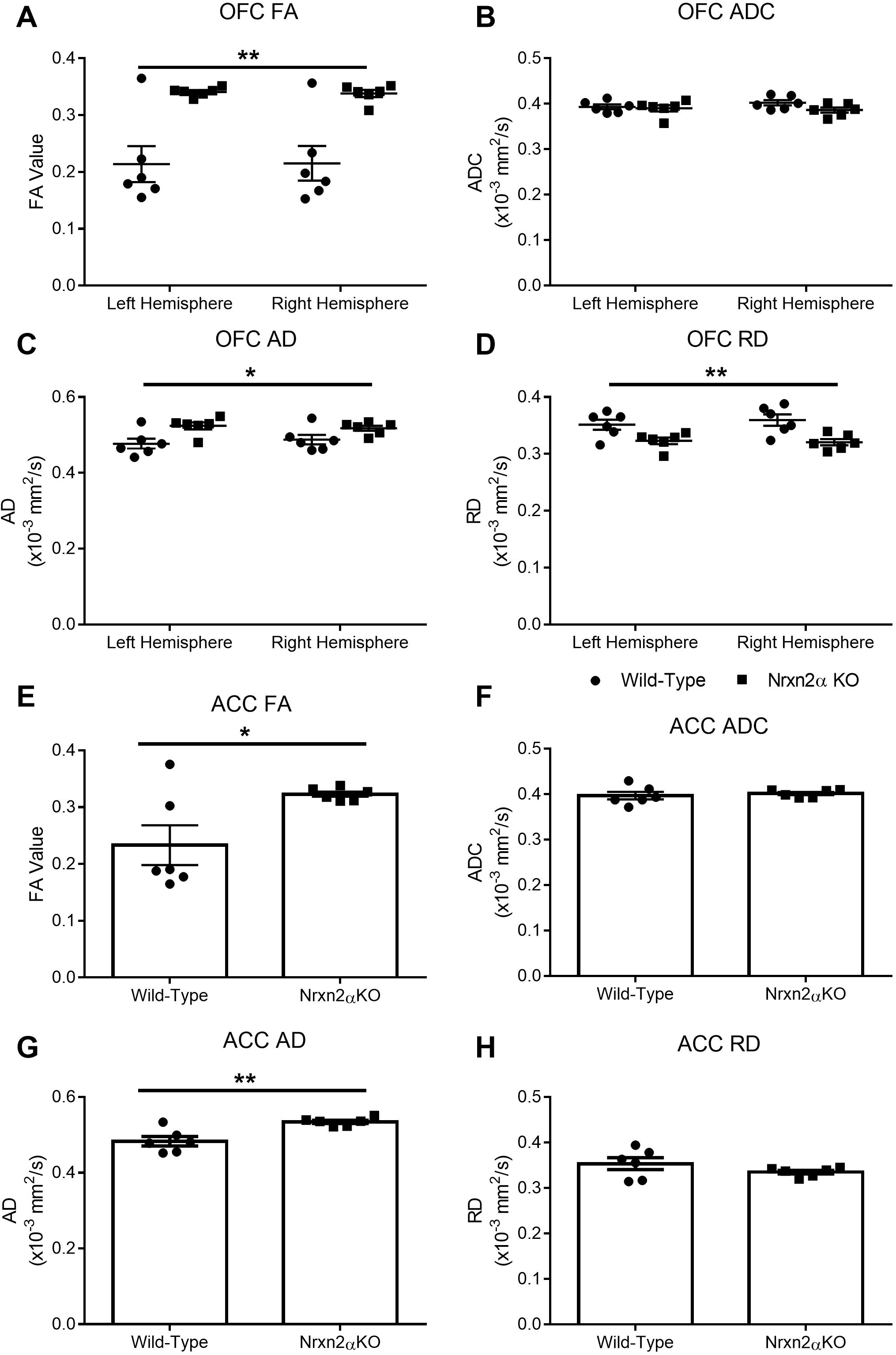
*Nrxn2α* KO mice have increased fractional anisotropy (FA) and axial (AD) and radial diffusivity (RD) in the orbitofrontal cortex (OFC) and the anterior cingulate cortex (ACC). FA was significant different between wild-types and *Nrxn2α* KO mice for FA in the OFC (**A**) and ACC (**E**), but ADC was not significantly changed in *Nrxn2α* KO mice in both prefrontal regions (**B** and **F**). The OFC has significantly increased AD and RD (**C&D**), whereas only AD was increased in the ACC (**G-H**). **=P<0.01, *=P<0.05. Error bars represent s.e.m. Wild-type n=6, *Nrxn2α* KO n=6.

We sought to examine whether changes in the amygdala, OFC or ACC FA and ADC were driven by diffusion in the primary axis (λ_1_) or the radial orientations (λ_2_ and λ_3_) by characterisation of AD (primary) and RD (radial). Within the amygdala, neither AD or RD was significantly altered in *Nrxn2α* KO mice (Figure 2E AD: genotype: F_(1,10)_ = 3.06, p = 0.111, Figure 2F RD: genotype: F_(1,10)_ = 2.47, p = 0.147). Within the OFC (Figure 3C&D), AD was significantly increased (genotype: (F_(1, 10)_ = 6.71, p = 0.032, power = 64.7%)), whereas RD was significantly decreased (genotype: (F_(1, 10)_ = 10.07, p = 0.025, power = 81.5%)), suggesting that both along-tract diffusion and tract branching were affected. However, in the ACC (Figure 3G-H), only AD was significantly increased (t_(10)_ = 3.89, p = 0.019, power = 96.9%), with no alteration in RD (t_(10)_ = 1.35, p = 0.10). Increased AD and decreased RD is thought to reflect changes in axonal density or orientation (54).

### DTI reveals altered hippocampal microstructure in *Nrxn2α* KO mice

The hippocampus has recently been associated with social motivation and social recognition. Since the specific contributions of the dorsal and ventral hippocampal poles remain unclear, we segmented the whole hippocampus into anterior (Bregma −1.06 mm – −2.46 mm) (incorporating dorsal) and posterior (Bregma −2.54 mm – −3.16 mm) (incorporating ventral regions) levels.

FA values in the anterior and posterior hippocampus were significantly increased (Supp. Figure 4A&E; see figure legend for statistics). However, ADC was unaltered for the anterior and posterior hippocampus (Supp. Figure 4B&F). AD was significantly increased in both the anterior and posterior hippocampal regions (Supp. Figure 4C&G). RD was significantly also significantly decreased in the anterior and posterior hippocampus in *Nrxn2α* KO mice (Supp. Figure 4D&H).

**Figure 4.**
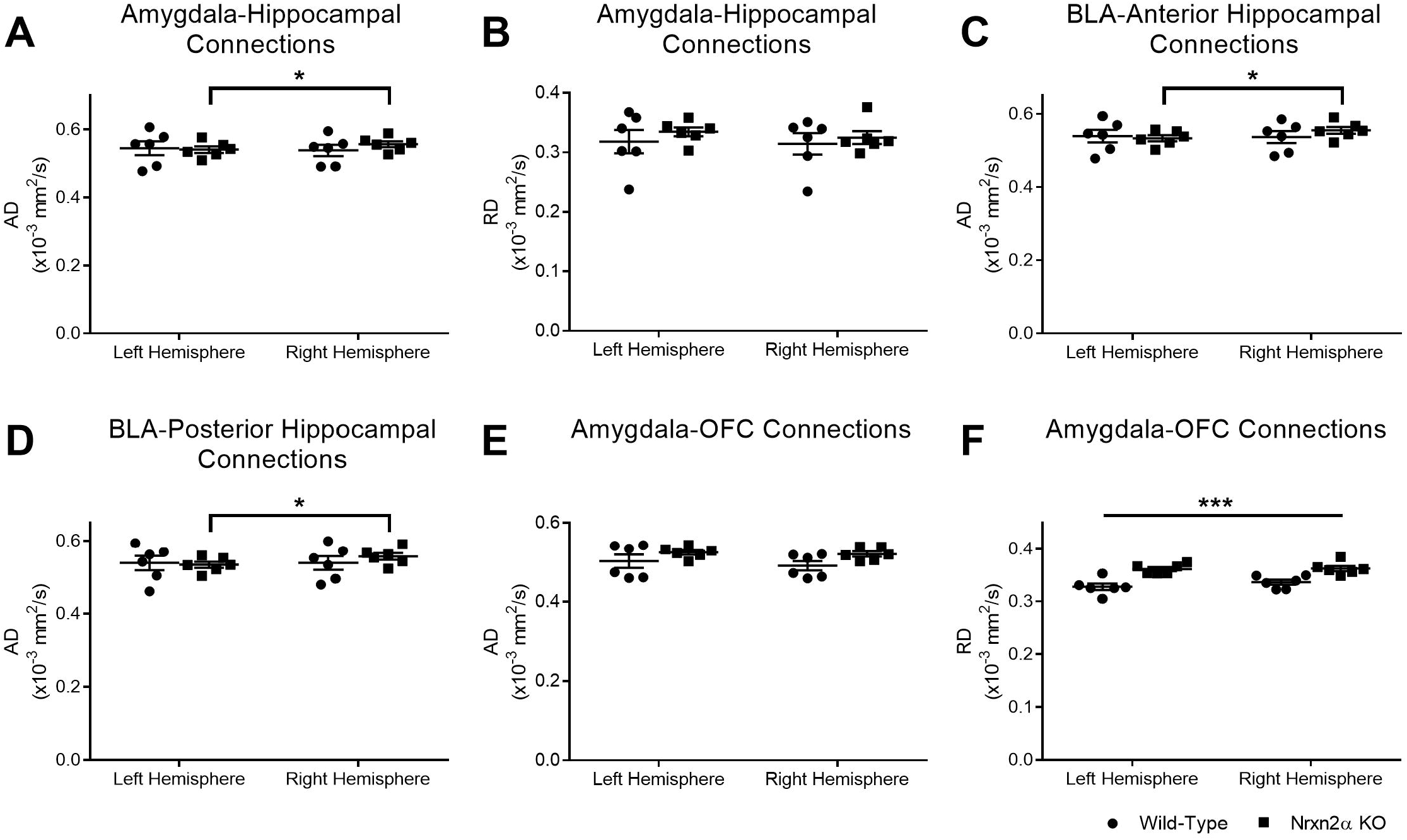
Tractographic analysis of amygdala-hippocampus and amygdala-orbitofrontal cortex (OFC) connectivity. Amygdala-hippocampal connections are characterised by greater right hemisphere axial diffusivity (AD) *Nrxn2α* KO mice (**A**) but not radial diffusivity (RD) (**B**). Specific to the BLA, connections to the anterior hippocampus (**C**) and posterior hippocampus (**D**) have greater right hemisphere AD. Although the amygdala-OFC connection was similar between the genotypes for AD (E), *Nrxn2α* KO mice had significantly increased RD (**F**). *=P<0.05, ***=P<0.001. Error bars represent s.e.m. Wild-type n=6, *Nrxn2α* KO n=6.

Lastly, given DTI is most commonly associated with analysis of white matter tracts, we also quantified the corpus callosum. Changes within the corpus callosum have repeatedly been highlighted in autism (55, 56), including mouse models of autism (57, 58). Here, we found significantly increased FA and reduced ADC in *Nrxn2α* KO mice, which were driven by a significant reduction in RD (Supp. Figure 6).

In summary, the microstructural measures most altered by *Nrxn2α* deletion were increases in FA, AD and RD, including in the hippocampus, in line with recent work suggesting a role for ventral hippocampus in social memory (43).

### DTI tractography reveals *Nrxn2α* deletion affects strucutral connectivity between the amygdala and orbitofrontal cortex

The amygdala is strongly and bidirectionally connected to both the hippocampus (59) and the OFC (60). As all three regions are themselves important for social behaviour, and autism is thought to be, at least in part, related to abnormal structural connectivity (24), we performed tractography analysis between the amygdala (and specifically the BLA) and the hippocampus, and between the amygdala and the OFC.

From the anterior amygdala, we examined the diffusivity (AD and RD) of connections to the anterior and posterior hippocampus (Supp. Figure 6). We did not observe differences in RD in the tracts connecting the amygdala with the hippocampus (see Supp. Table 1 for non-significant statistics). Although AD between the anterior amygdala and anterior hippocampus did not differ by genotype, there was a significant interaction between genotype and hemisphere (genotype × hemisphere (F_(1, 10)_ = 12.12, p = 0.023, power = 88.0%; Figure 4A); post hoc analysis shows this was driven by larger right-vs-left hemisphere AD values within the *Nrxn2α* KOs only (p = 0.012). This difference could be driven by the BLA; there was increased AD in both the BLA/anterior hippocampus tracts (genotype × hemisphere (F_(1, 10)_ = 10.53, p = 0.032, power = 83.2%) and the BLA/posterior hippocampus tracts (genotype × hemisphere (F_(1, 10)_ = 12.97, p = 0.020, power = 90%), which again was related to larger right-vs-left hemisphere values in the *Nrxn2α* KOs (BLA/anterior hippocampus: p = 0.004 and BLA/posterior hippocampus: p = 0.001, (Figure 4C-D)) but not the wild-type (anterior: p = 0.87; posterior: p = 1.00). These results indicate that there are differences for the structural connectivity of the amygdala with the hippocampus within the left and right hemisphere in *Nrxn2α* KO mice, with increased axial diffusivity in the right hemisphere. This finding is particularly interesting, as hemispheric differences in functional connectivity, particularly affecting connections from the right amygdala, have been found children with ASD (61, 62).

Finally, we tested connections between the amygdala and the OFC. For AD, wild-type and *Nrxn2α* KO mice did not differ by genotype (Figure 4E: genotype: (F_(1, 10)_ = 2.85, p = 0.09), hemisphere: (F_(1, 10)_ = 6.38, p = 0.052). RD was strikingly higher in *Nrxn2α* KO mice (Figure 4F: genotype: (F_(1, 10)_ = 26.06, p = 0.023, power = 99.5%)), indicative of a change in demyelination, axonal density or orientation (54).

### CLARITY reveals fibre disruption in *Nrxn2α* KO mice in the amygdala, orbitofrontal cortex, and anterior cingulate cortex

To further explore the differences as revealed by DTI, we performed CLARITY on the same brain tissue used in DTI, and stained with neurofilament and DAPI to label axons and cell bodies, respectively. We were then able to derive both the axonal alignment (as in, the geometric alignment of axons (from linear alignment to random) within 3D space (see Supp. Figure 2)) and density of the stained fibers, in addition to the cell density.

The pattern of results was broadly similar for both the prefrontal cortical ROIs. That is, first, axonal alignment was increased in *Nrxn2α* KO mice in the ACC (Figure 5D: genotype: (F_(1, 10)_ = 16.06, p = 0.011, power = 94.9%) but not the OFC (Figure 5G: genotype: (F_(1, 10)_ = 5.56, p = 0.059). Second, this could not be explained by a difference in cell density, since that was similar between the KO and wild-type mice in both the ACC (Figure 5F: genotype: (F_(1, 10)_ <1), hemisphere: (F_(1, 10)_ = 1.73, p = 0.11) and the OFC (Figure 5H: genotype: (F_(1, 10)_ = 3.09, p = 0.08). An increase in axonal density in *Nrxn2α* KO mice was reliable in the ACC (Figure 5E: genotype: (F_(1, 10)_ = 14.64, p = 0.014, power = 93.0%), but not in the OFC (Figure 5H: genotype: (F_(1, 10)_ = 3.09, p = 0.083).

**Figure 5.**
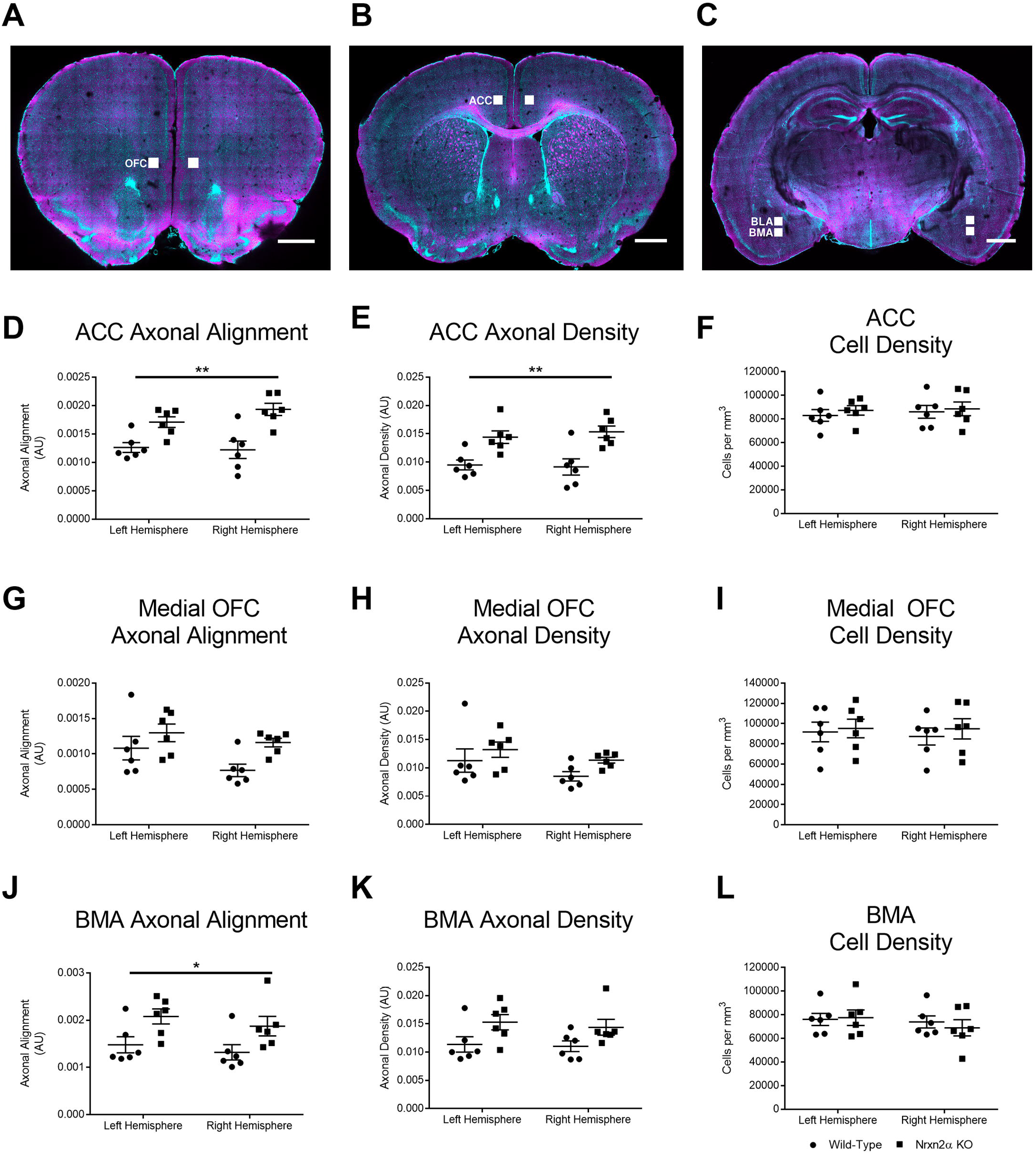
CLARITY reveals differences in axonal alignment and fibre density in *Nrxn2α* KO mice. (**A-C**) Representative images of the CLARITY-treated brain, with ROI defined for the anterior cingulate cortex (ACC), orbitofrontal cortex (OFC), basomedial amygdala (BMA) and basolateral amygdala (BLA). For the ACC, the axonal alignment (**D**) and axon density (**E**) were significantly altered in KO mice, but cell density was unaltered (**F**). Within the medial OFC, only axonal alignment was significantly altered in KOs (**G**), with axon density (**H**) and cell density (**I**) being similar. For the BMA, both the axonal alignment (**J**) and axon density (**K**) were significantly increased, whilst cell density was unaltered (**L**). *=P<0.05, **=P<0.01. Error bars represent s.e.m. Wild-type n=6, *Nrxn2α* KO n=6.

We further examined two regions of the anterior amygdala, the BLA and basomedial (BMA) nuclei, where altered social cellular responses have been reported in human autism (38). We did not observe any significant differences for axonal alignment or fibre density in the BLA (see Supp. Figure 7A-C), but whereas axonal alignment (Figure 5J, genotype: F_(1, 10)_ = 7.70, p = 0.045, power = 70.6%) but not axonal density (Figure 5K: genotype: (F_(1, 10)_ = 6.10, p = 0.054) was increased in *Nrxn2α* KO mice in the basomedial nuclei, while cell density was unaffected (Figure 5L: genotype: (F_(1, 10)_ <1). Alterations in axonal alignment and density as directly revealed by CLARITY could explain the increases in diffusivity and RD in the prefrontal regions, as measured by DTI.

To test the specificity of these alterations, we examined three further brain regions; the primary motor cortex (M1; Supp. Figure 7D-F), the primary somatosensory cortex (S1; Supp. Figure 7H-J) and the barrel field (BF; Supp. Figure 7K-M). Interestingly, although there were differences between the hemispheres, there were no statistical differences between the genotypes or genotype × hemisphere interactions for any measure (Supp. Table 2), suggesting some specificity of the alterations in social-relevant brain regions in *Nrxn2α* KO mice.

In summary, in both the prefrontal ROIs, namely the OFC, and the ACC, DTI showed that ADC and RD were increased in *Nrxn2α* KO mice, likely related to complementary analysis from CLARITY showing that axonal alignment was altered in *Nrxn2α* KO mice in both prefrontal ROIs.

## Discussion

Interestingly, the single-gene deletion of *Nrxn2α* captures several key aspects of human ASD. In terms of behaviour, three studies have now found social deficits associated with *Nrxn2α* KO (17-19); in terms of brain structure, as reported here (summarised below), the *Nrxn2α* KO mouse model shows altered microstructure and structural connectivity patterns in socially-relevant brain regions reminiscent of changes in ASD.

A DTI approach has been used for some time to explore neuropathological markers in autistic patients; alterations in myelination, axonal abundance, size and orientation all modify FA and ADC values (21, 63), specifically by reducing amygdala FA (23, 63), and have been used as a quantitative measure of changes to brain white matter integrity (23, 24). However, several studies have noted increases in FA in ASD patients (see Table 1 of 64). Furthermore, both increased RD of various white matter tracts (65, 66) and increased whole-brain AD (66) have been observed in ASD. The *Nrxn2α* KO mouse reproduces some of these specific changes, including altered FA and increases in ADC, AD and RD. Whole brain increases in ADC, AD and RD (but not FA) have been reported in ASD children, as have increases in ADC and RD in frontal cortex tracts (66). FA has been noted as reduced in the amygdala in ASD children and adolescents (67), and right-sided lateralization of abnormal amygdala/hippocampus-related connections, as seen in our *Nrxn2α* KO mouse, has been noted in high-functioning adolescents/adults with autism (68).

Whilst the current study specifically explores structural connectivity, it is difficult to extrapolate as to what these structural changes mean for functional connectivity in the *Nrxn2α* KO mouse. Hyper and hypo connectivity theories of autism have remained contentious, and vary in humans by cohort studied (e.g. by age of participant) (69). Further, in studies that have combined resting state functional MRI (rsfMRI) and DTI, functional and structural connectivity do not always overlap (70-72). Our current data suggests that DTI differences can be explained by altered axonal patterning (i.e. CLARITY). Others have explored the biological mechanisms linking structural connectivity to altered functional connectivity. Zhan et al. (2014) found that deletion of the chemokine receptor Cx3cr1 resulted in impaired synaptic pruning of long-range connections during development, which manifested as impaired social behavior caused by decreased frontal functional connectivity, reduced synaptic multiplicity and weakened coherence of local field potentials (73). Thus, it is possible that impairments in neuronal structural maturation can generate functional connectivity deficits that encapsulate core autism phenotypes.

Our findings corroborate these quantifications of clinical autism, but highlights the question of what do the different measures of ADC, FA, AD and RD represent? Importantly, we observed these microstructural changes in various socially-relevant brain regions against a background of unchanged cell density in all our study’s ROIs. Unexpectedly, this highlights the power of our new approach. Dudanova et al. (2007) concluded from measures of cell counting and cortical cell layer thickness that *Nrxn2* played little role in normal brain development (20). Indeed, in earlier studies, it was suggested that deletion of all *Nrxns* was unlikely to affect synaptic development but instead disrupts synaptic function (74). We propose that measures such as two-dimensional cell counting may be underestimating the impact of genetic mutations upon normal development. By staining cleared brain tissue with a nuclear marker and performing automated three-dimensional cell counting, we found no effect of *Nrxn2α* deletion on cell density in any region of interest examined. But this belies the clear effects upon microstructure integrity across multiple regions as measured by both DTI and CLARITY, and its specificity; only the socially-relevant brain regions we tested were disrupted, and not primary sensory or motor regions. Future studies will benefit from employing more sensitive measures of brain structural connectivity to determine the relevance of genetic mutations in development.

FA and ADC can be influenced by changes in axonal density and alignment (e.g. by myelination, demyelination, axonal damage, loss of white matter coherence (75)). It is likely that the axonal alignment metric used to quantify CLARITY more closely reflects the ADC measure of DTI, given that ADC (or mean diffusivity) equally weights diffusion across all eigenvectors and does not bias the primary eigenvector as FA does. Thus, it is likely that alterations in the properties of axons in *Nrxn2α* KO mice are driving these changes in FA and ADC. Given we see differences in RD, thought to reflect tract branching and myelination (as it measures λ_2_ and λ_3_), it is possible that the orientation in the perpendicular not parallel orientation of fibres is mostly affected. Given the differences in the amygdala, OFC and ACC, it is possible that even though neuronal densities are similar in the *Nrxn2α* KO brain, it is the connections between neurones and brain regions that are perturbed. This would be consistent with the idea that structural connectivity disruption may represent a core feature of autism (76). A broader question is how does the loss of *Nrxn2α* account for changes in axonal organisation? Ultimately, this question requires further studies. Others have shown that in *Nrxn2α* KO mice, excitatory transmitter release is reduced, as is short-term plasticity (18). Reduced glutamatergic release, even at a relatively long range to the synapse, can change the complexity of dendritic arbors (77). As this is a gene deletion model, it is conceivable that altered glutamatergic signalling during early development impairs appropriate synapse maturation, leading to the structural changes we see herein. Further, how or whether these structural changes fully explain the social impairments of *Nrxn2α* KO mice would require new studies. Conceivably, inducible knock-down of *Nrxn2* (by inducible knockout, siRNA, optogenetics etc.) within a specific brain region would provide evidence that social abnormalities are being driven by *Nrxn2* loss. However, developmentally-dependent altered structural connectivity would be harder to definitively manipulate to explain changes in social behaviours.

Here we have developed a new application of CLARITY to quantitatively investigate disease models by combining DTI with high resolution 3D imaging and automated analysis of axonal fibres in a within subject study. Inevitably, there are some technical limitations that will require future refinement as this technology matures.

First, while we used CLARITY and immunolabeling to identify axons, we cannot know whether axon-related changes alone reflect all the changes we observed for our DTI measures. Second, whilst we can segment entire brain regions for DTI analysis, it was not practical to image larger brain areas at the necessary resolution for CLARITY. While it is theoretically possible that we may bias sampling of each brain region by picking ROIs for multiphoton imaging, this was done using atlas-defined coordinates and by an experimenter blind to the DTI results, so minimising any bias. However, within the current study, we were only able to apply the CLARITY approach to the amygdala, OFC and ACC. It was not practical to apply this methodology to the hippocampus, due to its extremely heterogeneous structure. The small cubic ROIs could not be reproducibly positioned, and larger ROIs to average across larger areas of the hippocampus were not possible. Although imaging of fibre tracts in large volumes of cleared tissue is possible (78), fluorescent labelling limitations make it impractical for a study of this nature. Despite this, as the adoption of the CLARITY technique increases, we hope that the use of DTI and CLARITY to study structural connectivity across spatial scales will become commonplace.

As yet, no one DTI protocol has emerged as the standard for *in vivo* or *ex vivo* imaging. Indeed, there has been debate regarding the best number of diffusion gradients to use, among other parameters (79). Undoubtedly, more directions that what we used here would facilitate better interpretations, and this is a limitation of the current work. Despite this, the major purpose of the current paper is to develop a new generation of CLARITY analysis. We hope that future studies will refine on both DTI and CLARITY parameters to maximise analysis methodology. A further potential limitation of the current study is that groups of six animals may be underpowered. We argue for our approach here as follows. First, low variance in the datasets permits smaller group sizes. Second, for most of our significant results, the observed power was more than 80%. Third, given the technical complexity of this approach, particularly in its early adoption and refinement stages, large sample throughput of multiple brain regions is challenging.

In summary, our combined use of DTI and CLARITY has revealed changes in microstructure and structural connectivity of socially-relevant brain regions in *Nrxn2α* KO mice that may underlie their deficits in social behaviour. It is hard to conceive how these changes could have been observed using classical experimental approaches. We envisage this approach will deliver a new level of detail in structural connectivity approaches to understanding autism.

## Abbreviations

ACC: anterior cingulate cortex
AD: axial diffusivity
ADC: apparent diffusion coefficient
ASD: autism spectrum disorder
BLA: basolateral amygdala
CLARITY: optically cleared brain tissue
DTI: diffusion tensor imaging
FA: fractional anisotropy
OFC: orbitofrontal cortex
Nrxn2: neurexin II
RD: radial diffusivity
ROI: region of interest

## Declarations

### Ethics approval and consent to participate

All experiments were performed under UK Home Office Project and Personal Licenses in accordance with the Animals (Scientific Procedures) Act 1986, and with the approval of the University of Leeds and Durham University Animal Ethical and Welfare Review Boards.

### Consent for publication

Not applicable

### Availability of data and material

The codes used to quantify the CLARITY datasets are made available by author LCA by email request to authors LCA or JD, subject to reference to the current paper. The datasets used and/or analysed during the current study are available from the corresponding author on reasonable request.

### Authors’ contribution

EP, ALT, LCA and JD conceived the study. EP and ALT performed the experiments. EP, ALT, LCA and JD analysed the data. SJC, RJR, LCA and JD funded the study. All authors contributed to writing the paper. All authors have read and approved the final manuscript.

## Acknowledgements

Not applicable.

## Supplemental Materials and Methods

### Diffusion Tensor MRI Data Acquisition

Brain MR imaging was performed on a vertical 9.4 Tesla spectrometer (Bruker AVANCE II NMR, Ettlingen, Germany) with an 89 mm wide bore, 3 radio frequency channels with digital broadband frequency synthesis (6-620 MHz) and an imaging coil with diameter of 25 mm for hydrogen (1H). 3D images for each brain were obtained using a DT-MRI protocol (TE: 35 ms, TR: 700 ms, 10 signal averages). The field of view was set at 128 × 128 × 128, with a cubic resolution of 100 μm/pixel and a b value of 1200 s/mm^2^. For each brain, diffusion weighted images were obtained in 6 directions, based upon recent published protocols (80-84). The subject of the number of diffusions gradients has been debated (79), with studies suggesting limited benefits of using more than 6 directions in biological tissue (85-87). The imaging time for each brain was 60 hours.

### CLARITY Solutions

#### Hydrogel solution

2% PFA 2% acrylamide 0.05% bis-acrylamide and 0.25% VA-044 thermal initiator (2,2’-Azobis[2-(2-imidazolin-2-yl) propane] dihydrochloride) in PBS, pH 7.4.

#### Clearing buffer

8% Sodium dodecyl sulfate in 200mM boric acid, pH 8.5.

### Multiphoton imaging – methodological outline

Cleared samples were mounted in custom 3D printed chambers for two-photon imaging. Images were acquired using ZEN Black (Zeiss, Germany). DAPI signal was detected using a 485 nm short pass filter, and neurofilament using a 500-550 nm band pass filter. The power of the excitation laser was varied to maximise the dynamic range for each image, but all other parameters were kept constant. The images were analysed using custom MATLAB (version 9.1, The Mathworks Inc.) scripts. Two-dimensional images were visualised using ImageJ (88) and three-dimensional images using Vaa3D (89).

## Multiphoton imaging and analysis – image analysis method

### Pre-processing

Image files were loaded into MATLAB (The Mathworks Ltd.) using the BioFormats toolbox (90), and the raw image data were obtained along with the precise voxel dimensions from the metadata. Each two-dimensional (2D) image from the three-dimensional (3D) stack was initially corrected for uneven background illumination by element-wise division by a 2D reference image. This reference image was calculated as the mean 2D image through the 3D stack, which was smoothed using a 2D Gaussian kernel with a full-width at half maximum (FWHM) of 20 % of the geometric mean of the dimensions of the 2D image (mean dimension). The image was denoised by filtering the image with a Gaussian kernel with a standard deviation of one pixel. Background subtraction was carried out by subtracting a smoothed, filtered image (FWHM 10 % of the mean dimension). Each pixel was then smoothed using a 3D Gaussian kernel with FWHM of 1.5 μm (the largest axonal diameter expected according to Perge et al. (91).

### Segmentation

The numerical gradient of the image in each dimension (Δ*X*, Δ*Y*, Δ*Z*) was calculated, and these were combined to calculate the magnitude of the gradient 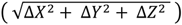. The resulting image was thresholded, using a combination of the Otsu (1979) and Rosin (2001) methods (Rosin threshold + 2/5 Otsu threshold) (92, 93). The gradient image highlights the edge of each axon; to combine these into a single object, the image was dilated and then eroded with a cubic structuring element (each side being 1.5 μm, to ‘close’ the largest axons as per Perge et al. (91)). Very small objects (less than 50 μm^3^) were removed from the image as they reflected noise, very small neuronal processes and gradients around cells.

Owing to variations in staining intensity of different axons, the thresholding produced segmented axons of various thicknesses that did not necessarily reflect the true structure. To remove this bias, the thresholded image was skeletonised using a homotopic thinning algorithm (94) implemented in MATLAB(95). The resulting image was dilated and then eroded using a cubic structuring element (10 pixels on each side for dilation, 9 for erosion) to produce connected processes with a uniform two-pixel diameter. This dilation ensured that the voxels in the binary image were connected via their faces (6-connected) rather than just their corners (26-connected), which better reflects the true structure of biological processes. This method detects most large axons at the expense of smaller processes, and the loss of any information about axon diameter. These steps are outlined in Supp. Figure 2.

### Analysis

The density of axons was calculated as the fraction of the image volume taken up by the segmented axons. A measure of axonal alignment was calculated by determining the mean axonal alignment along each dimension. This alignment was calculated by moving along the 3D image in a single dimension, keeping the coordinates in the other dimensions constant, and counting the number of times the pixel intensity did not change (i.e. how many times an axon was not entered or left). This number was averaged across each face of the image volume and scaled to the length of each dimension to produce a metric of how constant the image intensity is in that dimension. The perfect case of no intensity change (i.e. all axons are aligned perfectly with a particular dimension) gives a value of 1. The greater the difference between this measure in each three dimensions, the more aligned the axons must be (i.e. their directions are anisotropic). The standard deviation of this measure across the three dimensions was calculated as the axonal alignment.

The alignment calculation is illustrated in Supp. Figure 3 for a simple, two-dimensional case. Supp. Figure 3a shows the case of low axonal alignment, and Supp. Figure 3b shows the case of high axonal alignment. In each case, for illustration, each pixel represented by a small square on the grid is classed as either containing an axon or not. In the real images, the pixels are smaller, and are actually binary. In each axis, the number of pixel transitions in which the presence of an axon does not change is divided by the number of transitions, and the average is calculated. The standard deviation of this average for all axes is the measure of axonal alignment. When the alignment is low, the two averages are similar, and the standard deviation is low. When the alignment is high the two averages are very different, and the standard deviation is high. To analyse the real data, this same calculation is carried out in 3D, but in a much larger grid of voxels. Cell density was calculated as the number of cells per mm^3^.

**Supp. Table 1.**
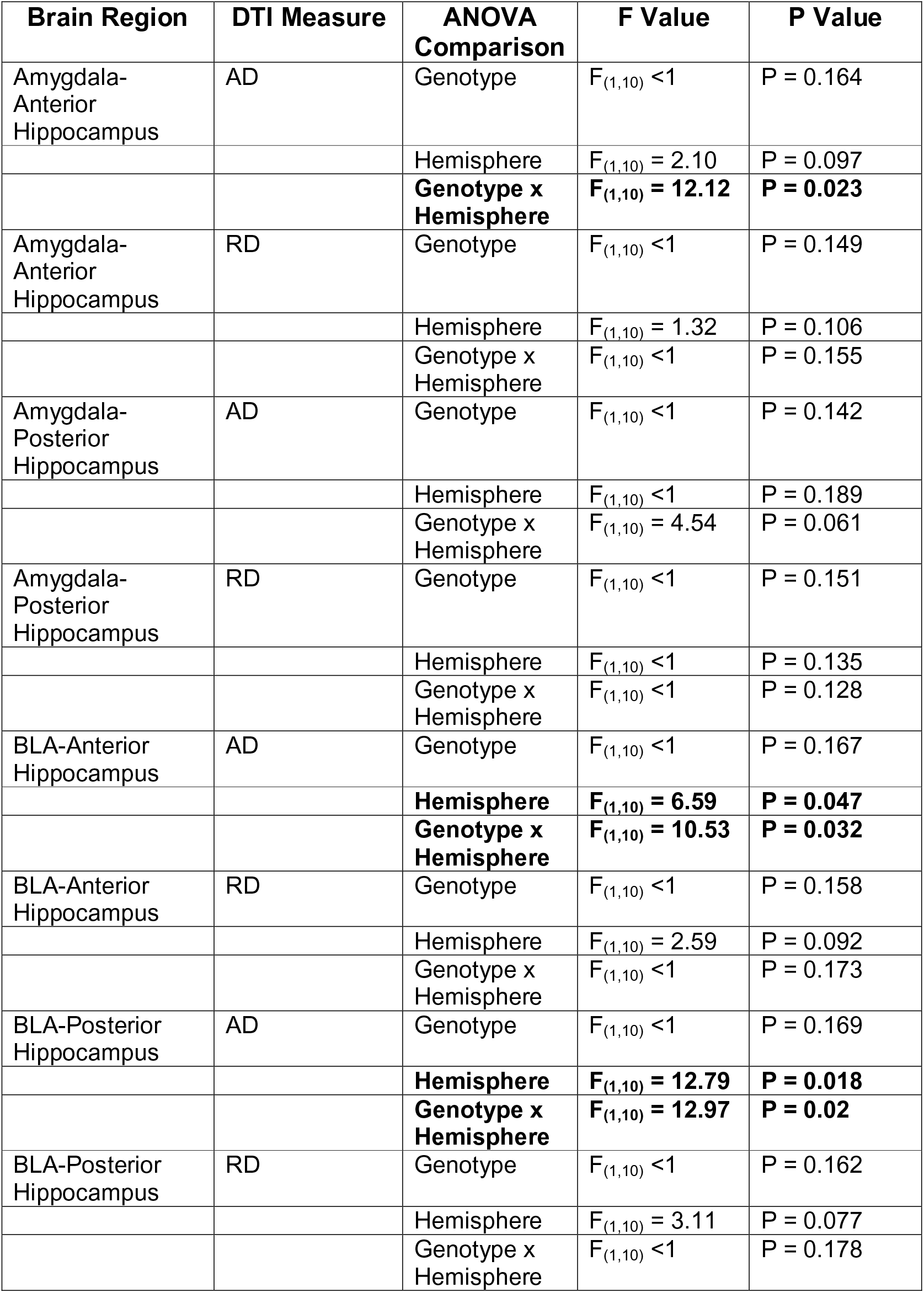
Statistical analysis of the anterior (Bregma −1.94 mm), and posterior (Bregma −3.28 mm) amygdala-hippocampal connections, analysed for axial diffusion (AD) and radial diffusion (RD). Analysis was performed using repeated measure two-way ANOVAs for genotype and hemisphere (Benjamini-Hochberg corrected (corrected P values stated)).

**Supp. Table 2.**
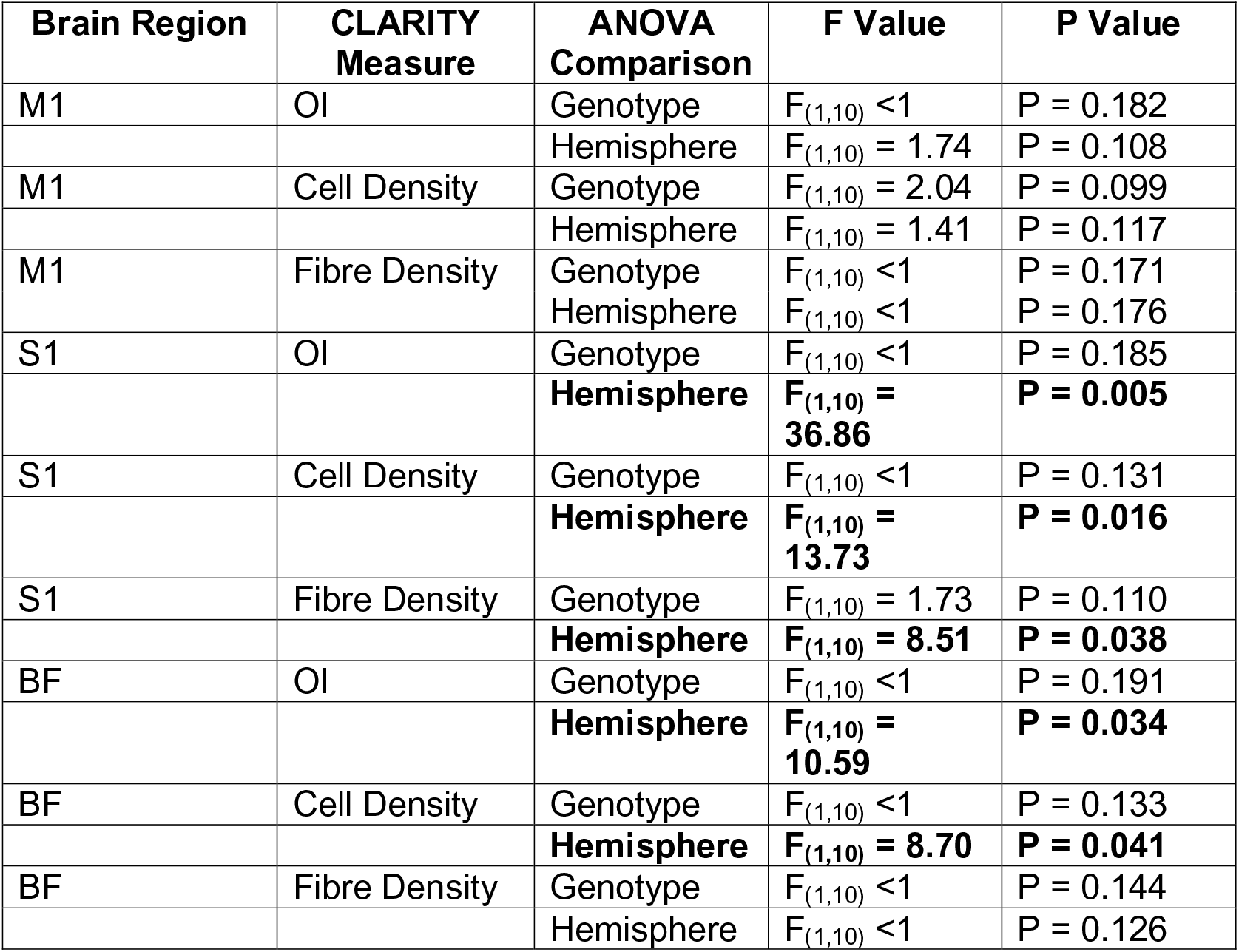
Statistical analysis of the primary motor cortex (M1), primary somatosensory cortex (S1) and the barrel field (BF). CLARITY imaged regions were then analysed for orientation index (OI), cell density and fibre density. Analysis was performed using repeated measure two-way ANOVAs for genotype and hemisphere (Benjamini-Hochberg corrected (corrected P values stated)).

**Supp. Figure 1.**
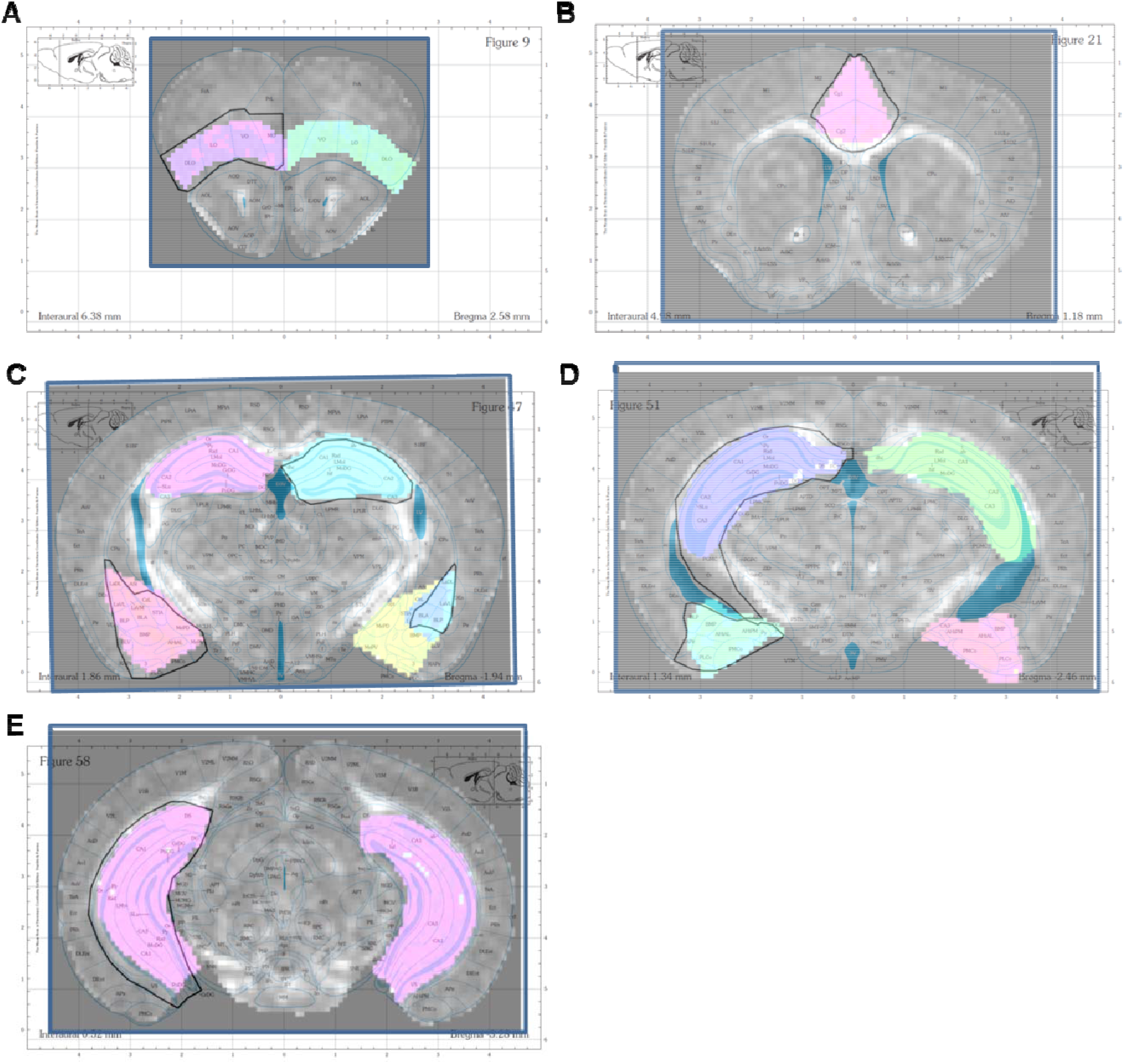
Atlas maps representing manual segmentation of regions of interest (ROI), overlaid with segmented brain regions from a fractional anisotropy-coloured brain slice. (**A**) The orbitofrontal cortex ROI. (**B**) The ACC ROI. (**C**) The anterior hippocampus, anterior amygdala and basolateral amygdala ROI. (**D**) The mid hippocampus and posterior amygdala ROI. (**E**) The posterior hippocampus ROI. The atlas maps were used with the permission of the Authors (96).

**Supp. Figure 2.**
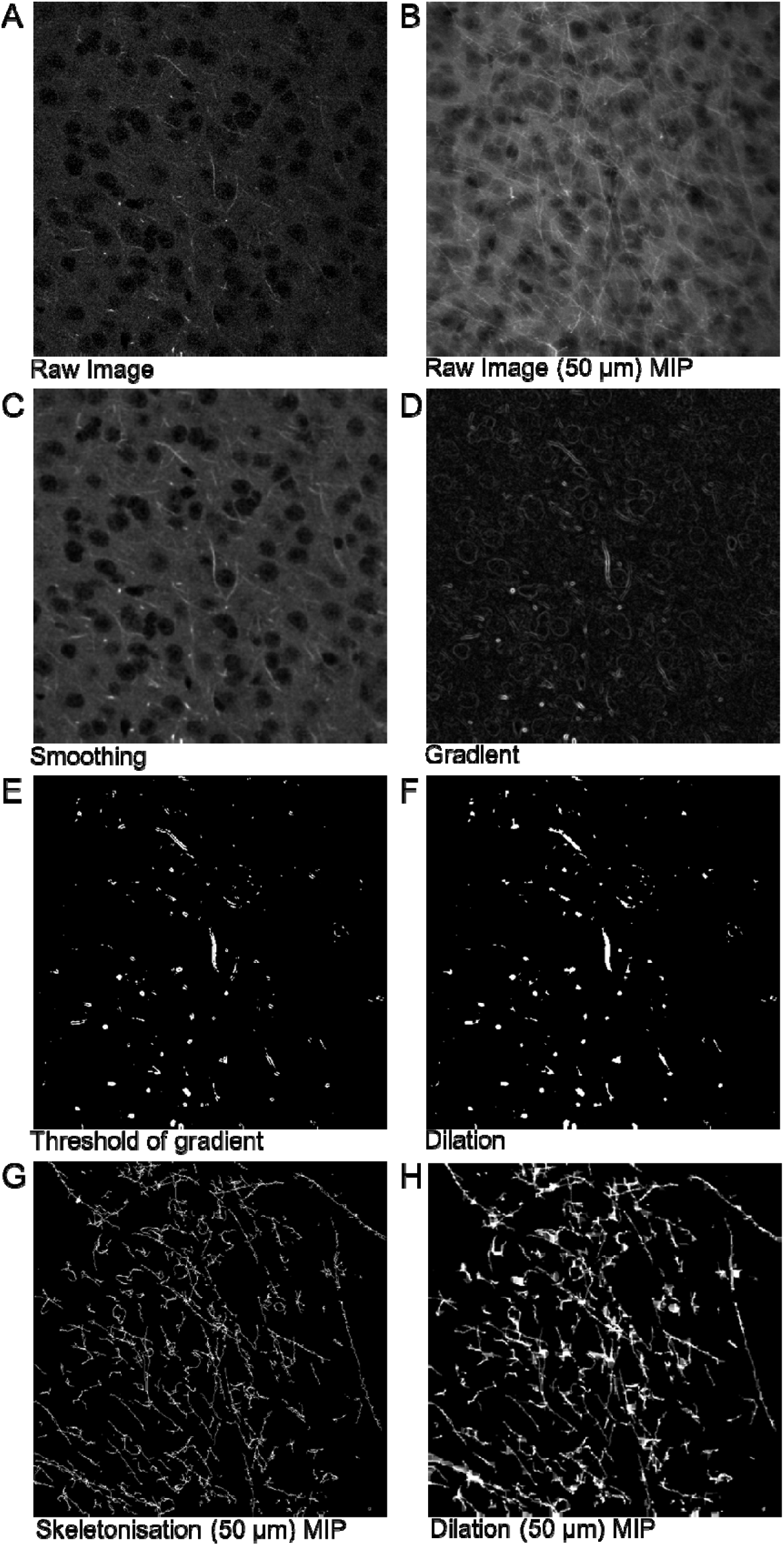
Analysis methodology of axonal segmentation from multiphoton images.

**Supp. Figure 3.**
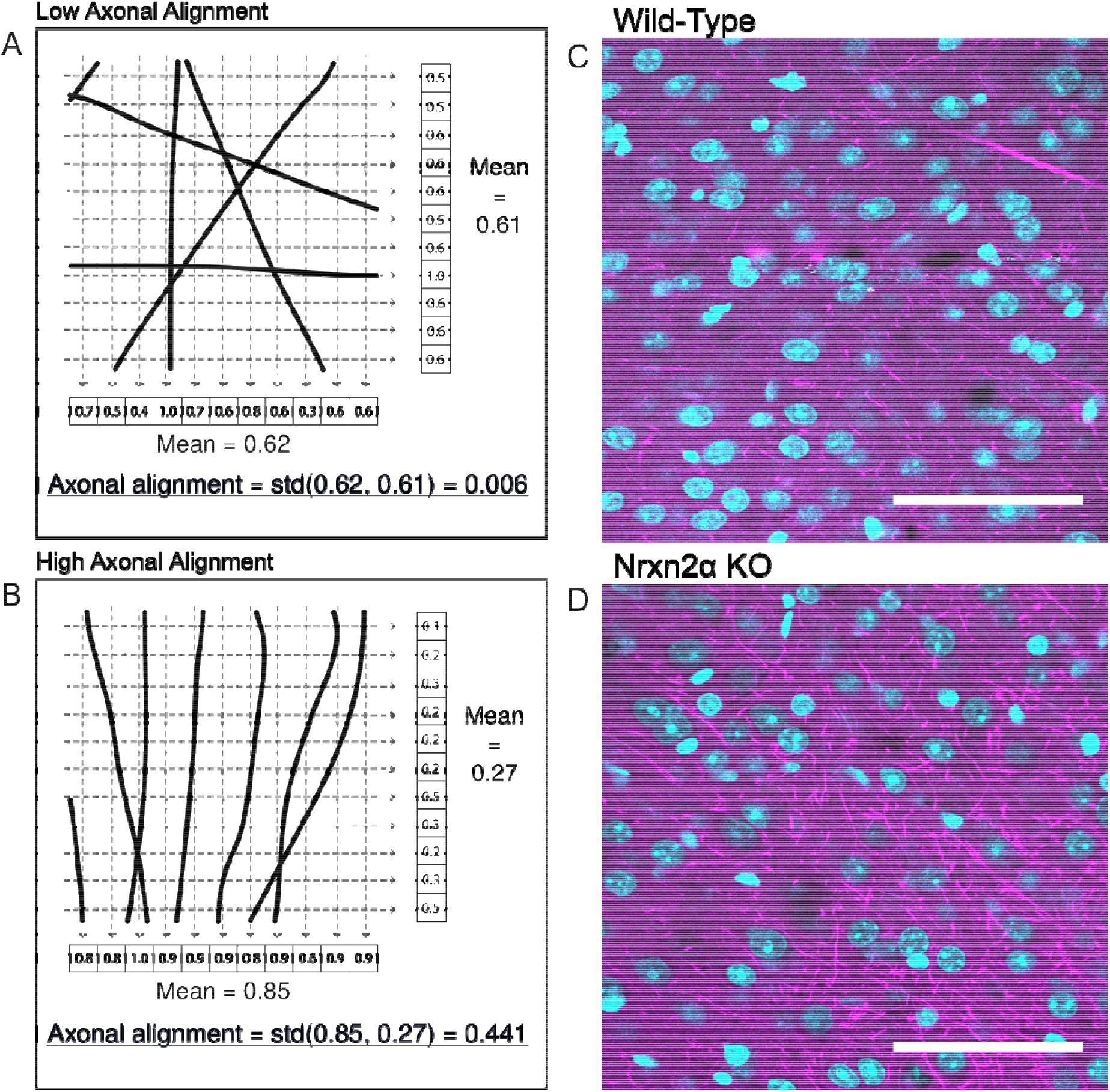
**A and B.** Illustration of the axonal alignment calculation in a simple two-dimensional case. The grey grid represents the image pixels, the black lines axons and the dashed lines represent the calculation process. Standard deviation denoted as std. **C and D.** Cingulate cortex images taken from wild-type and Nrxn2α KO mice, visually representing the greater axonal alignment and density in KO mice.

**Supp. Figure 4.**
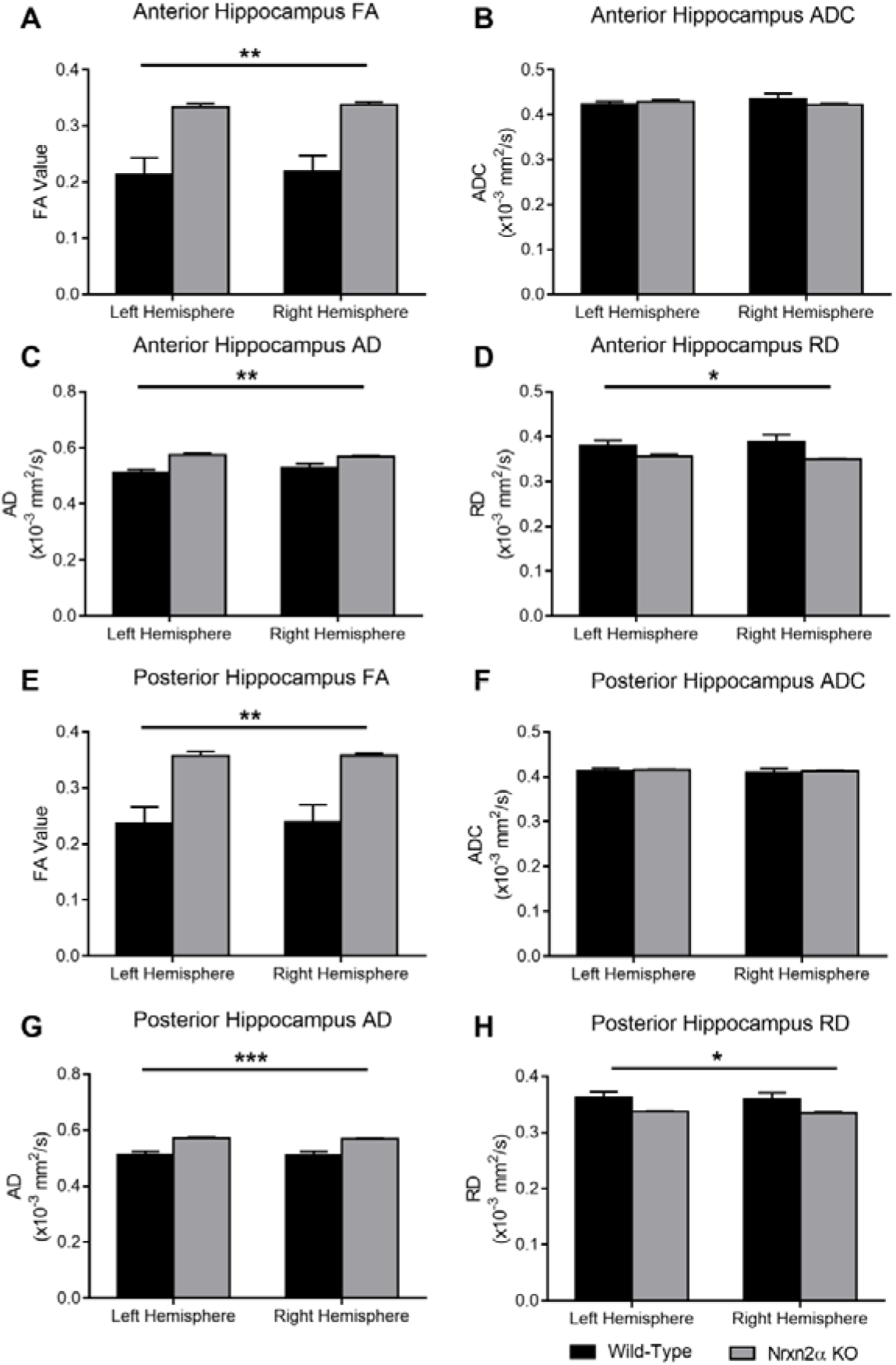
DTI quantified for the whole anterior hippocampus (Bregma −1.06 mm – −2.46 mm) and posterior hippocampus (Bregma −2.54 mm – −3.16 mm). (**A**) Fractional anisotropy (FA) in the anterior hippocampus was significantly increased in Nrxn2α KO mice (genotype: F_(1,10)_ = 15.63, p = 0.0027) but (**B**) apparent diffusion coefficient (ADC) was not altered (genotype: F_(1,10)_ <1, p = 0.738). (**C**) Axial diffusivity (AD) (genotype: F_(1,10)_ = 16.17, p = 0.0024) and (**D**) radial diffusivity (RD) (genotype: F_(1,10)_ = 5.05, p = 0.048) were both significantly altered in Nrxn2α KO mice. In the posterior hippocampus, in Nrxn2α KO mice, (**E**) FA was significant increased (genotype: F_(1,10)_ = 15.62, p = 0.0027), (**F**) ADC was similar to wild-types (genotype: F_(1,10)_ <1, p = 0.679), (**G**) AD was increased (genotype: F_(1,10)_ = 22.31, p = 0.0008) and (**H**) RD was significantly reduced (genotype: F_(1,10)_ = 5.34, p = 0.043). Error bars Nrxn2α KO n=6.

**Supp. Figure 5.**
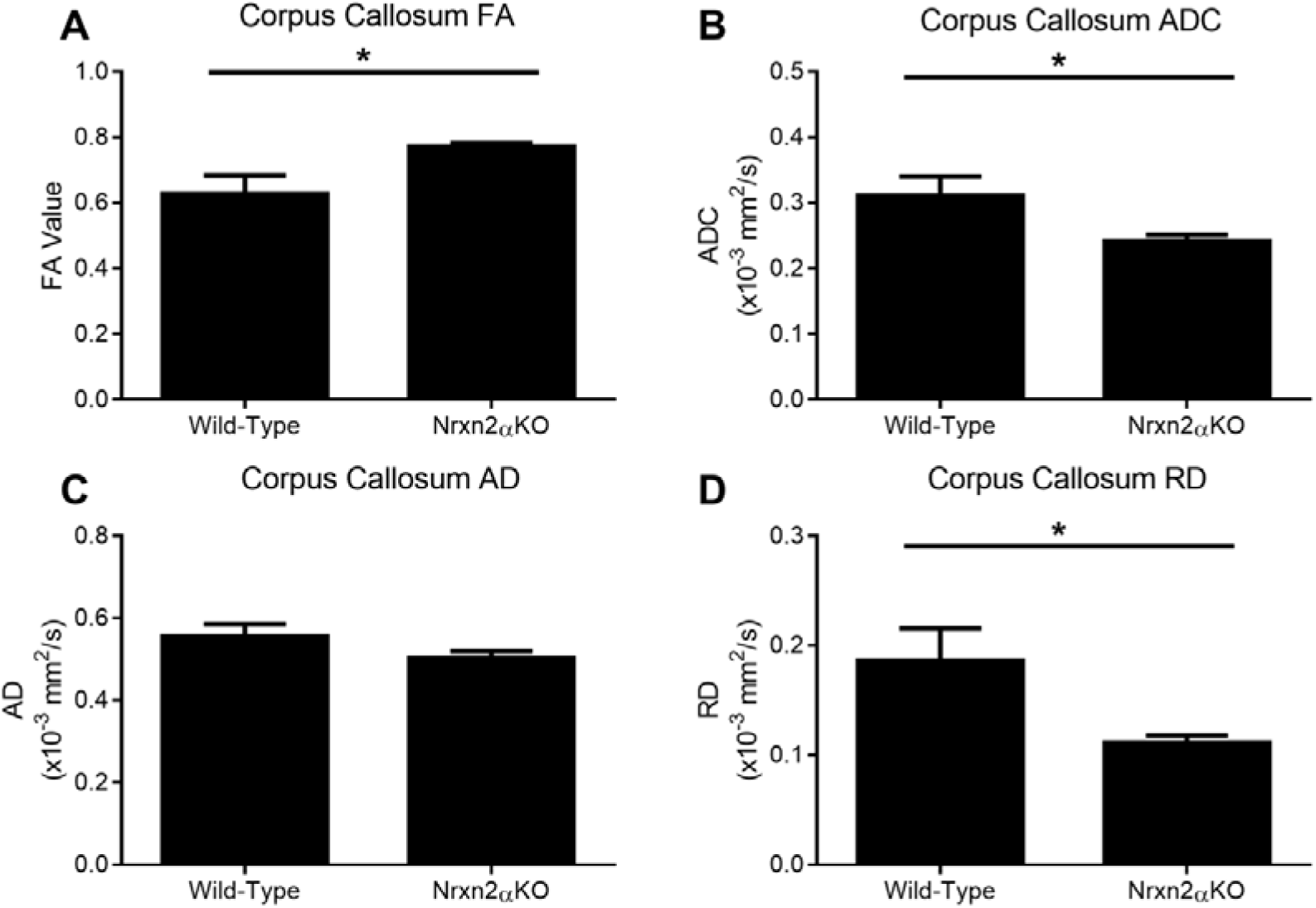
DTI quantification of the corpus callosum (Bregma 0.98 mm). To examine the integrity of white matter tracts within Nrxn2α KO mice, we examined diffusivity in the corpus callosum. (**A**) Fractional anisotropy (FA) was significantly increased in Nrxn2α KO mice (genotype: t_(10)_ = 2.50, p = 0.032) and apparent diffusion coefficient (ADC) (**B**) significantly decreased (genotype: t_(10)_ = 2.28, p = 0.046). This difference appeared to be driven predominantly by radial diffusivity (RD), as axial diffusion (AD) (**C**) was not significantly different (genotype: t_(10)_ = 1.49, p = 0.168) whilst RD (**D**) was significantly reduced in Nrxn2α KO mice (genotype: t_(10)_ = 2.45, p = 0.034). Error bars represent s.e.m. * = P<0.05. Wild-type n=6, Nrxn2α KO n=6.

**Supp. Figure 6.**
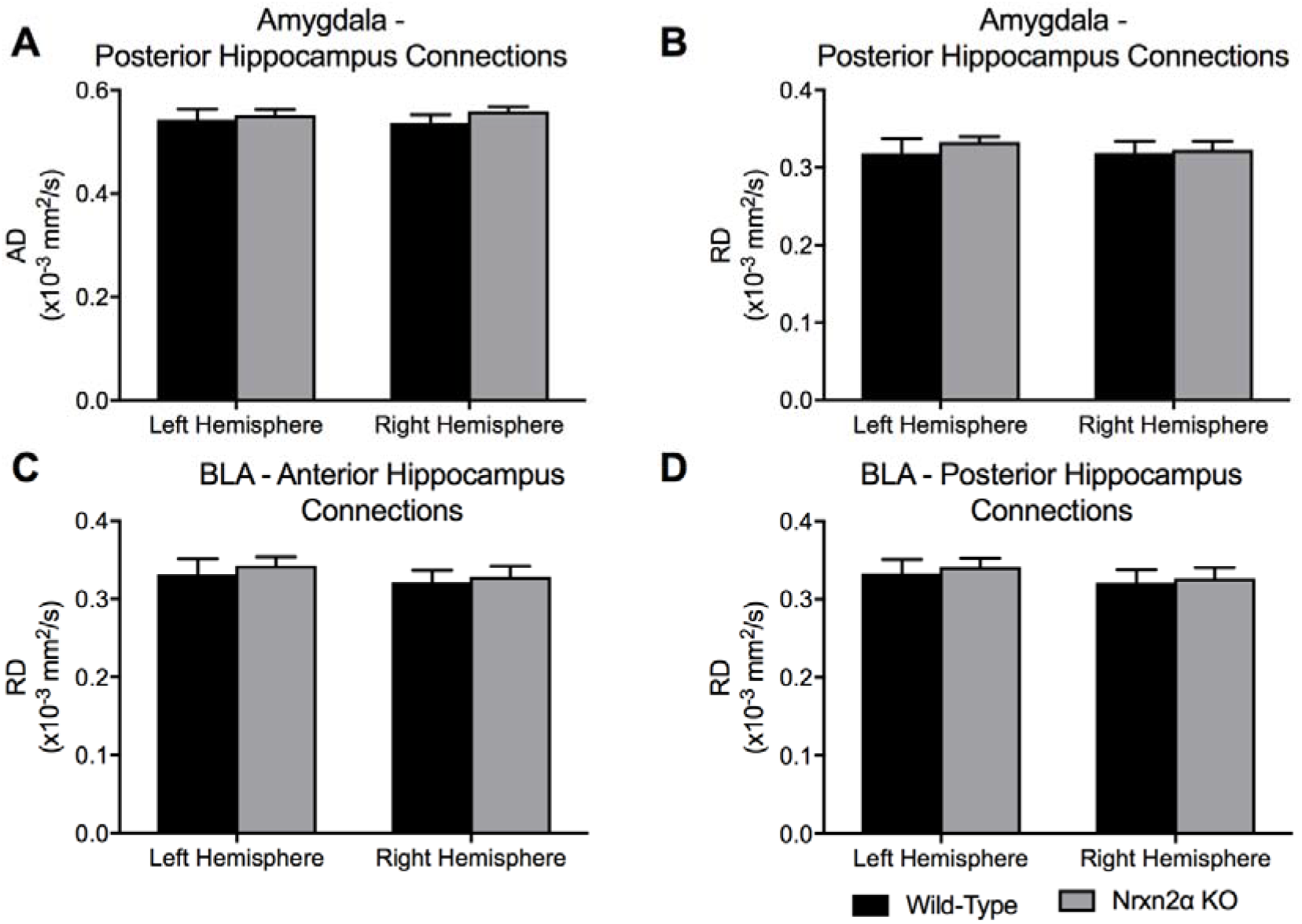
Axial diffusivity (AD) and radial diffusivity (RD) of computed tracts of connections from the amygdala to hippocampus. Tracts from the anterior amygdala to the posterior hippocampus (Bregma −2.46 mm) were analysed for AD (**A**) and RD (**B**). No significant differences between the tracts of Nrxn2α of KO mice were observed. No significant differences were found for RD of tracts specifically from the basolateral nuclei of the amygdala (BLA) to the anterior (**C**) or posterior (**D**) hippocampus. Error bars represent s.e.m. Wild-type n=6, Nrxn2α KO n=6.

**Supp. Figure 7.**
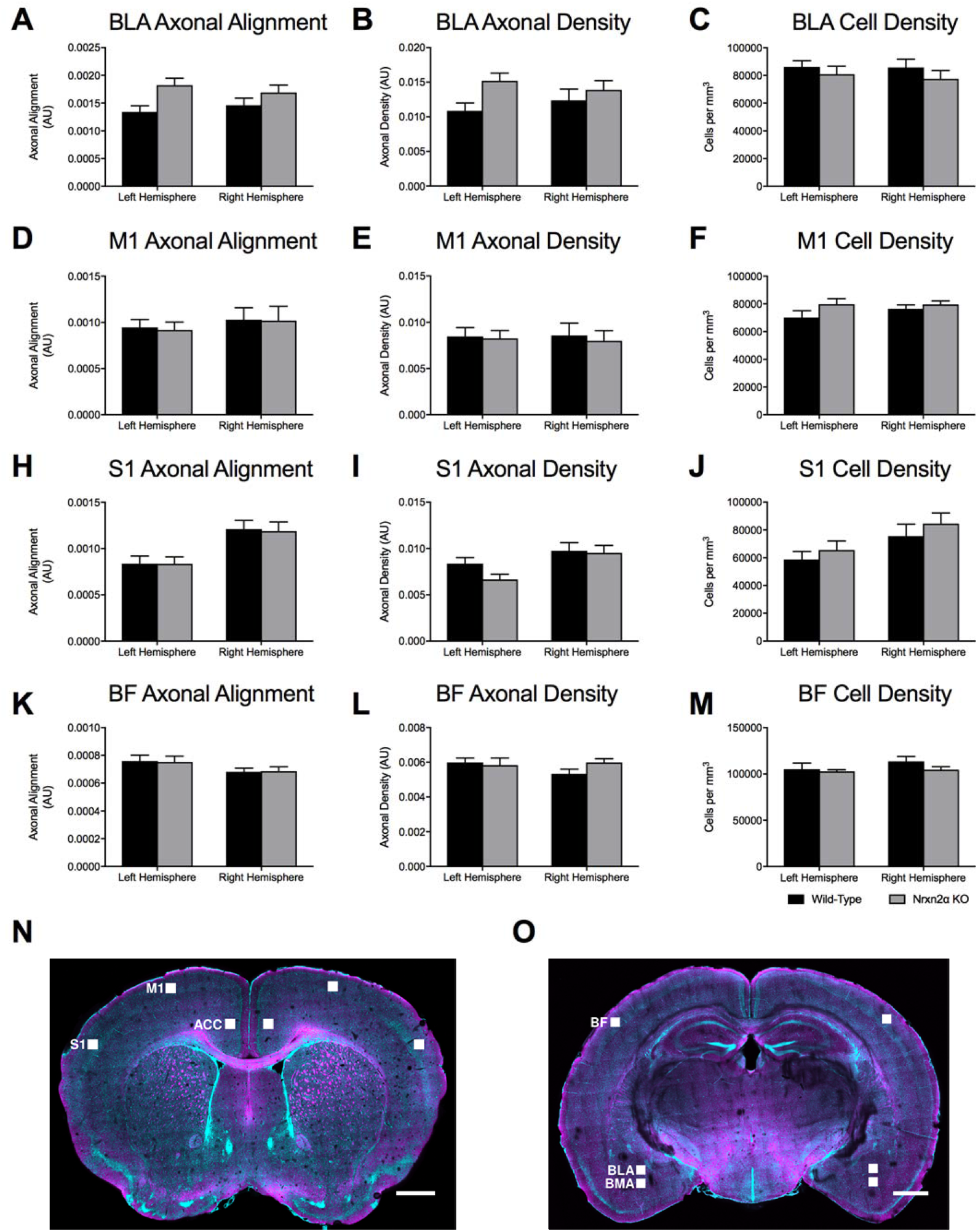
CLARITY-derived quantification of fibres and cell density within the basolateral amygdala (BLA) and control regions (**A**) Although there were trends towards increased axonal alignment and fibre density (B) in Nrxn2α KO mice, no significant differences were found. (**C**) Cell density in the BLA was similar between the genotypes. Statistical analysis (Supp. Table 4) was performed for the primary motor cortex (M1; **D-F**), primary somatosensory cortex (S1; **H-J**) and the barrel field (BF; **K-M**). No genotypic differences were found for any measure within these cortical regions. (N-O) CLARITY images of the scanned regions of interest. Error bars represent s.e.m. Wild-type n=6, Nrxn2α KO n=6.

