## Supplementary material for "The within-subject application of diffusion tensor MRI and CLARITY reveals brain structural changes in *Nrxn2* deletion mice": Supp Material

### Supplemental Material

Eleftheria Pervolaraki Ph.D, Adam L. Tyson MRes, Francesca Pibiri Ph.D, Steven L. Poulter Ph.D, Amy C. Reichelt Ph.D, R. John Rodgers Ph.D, Steven J. Clapcote Ph.D, Colin Lever Ph.D, Laura C. Andreae M.D. Ph.D, James Dachtler Ph.D

**Segmentation**

The numerical gradient of the image in each dimension $(\Delta X, \Delta Y, \Delta Z)$ was calculated, and these were combined to calculate the magnitude of the gradient $\left( \sqrt{\Delta X^2 + \Delta Y^2 + \Delta Z^2} \right)$. The resulting image was thresholded, using a combination of the Otsu (1979) and Rosin (2001) methods (Rosin threshold + 2/5 Otsu threshold) (14, 15). The gradient image highlights the edge of each axon; to combine these into a single object, the image was dilated and then eroded with a cubic structuring element (each side being 1.5 μm, to 'close' the largest axons as per Perge et al. (13)). Very small objects (less than 50 $\mu m^3$) were removed from the image as they reflected noise, very small neuronal processes and gradients around cells.

**Supp. Table 1**

| Brain Region | DTI Measure | ANOVA Comparison | F Value | P Value |
|---|---|---|---|---|
| Anterior Hippocampus | FA | Genotype | $F_{(1,10)}$ = 3.91 | P = 0.063 |
| | | Hemisphere | $F_{(1,10)}$ = 3.89 | P = 0.065 |
| Mid Hippocampus | FA | Genotype | $F_{(1,10)}$ = 3.12 | P = 0.07 |
| | | Hemisphere | $F_{(1,10)}$ <1 | P = 0.146 |
| Posterior Hippocampus | FA | Genotype | $F_{(1,10)}$ = 1.99 | P = 0.104 |
| | | Hemisphere | $F_{(1,10)}$ = 1.95 | P = 0.101 |
| Anterior Hippocampus | ADC | Genotype | $F_{(1,10)}$ = 3.17 | P = 0.073 |
| | | Hemisphere | $F_{(1,10)}$ = 2.20 | P = 0.095 |
| Mid Hippocampus | ADC | Genotype | $F_{(1,10)}$ <1 | P = 0.160 |
| | | Hemisphere | $F_{(1,10)}$ = 1.62 | P = 0.115 |
| Posterior Hippocampus | ADC | **Genotype** | **$F_{(1,10)}$ = 8.88** | **P = 0.036** |
| | | Hemisphere | $F_{(1,10)}$ <1 | P = 0.180 |
| Anterior Hippocampus | AD | Genotype | $F_{(1,10)}$ <1 | P = 0.187 |
| | | Hemisphere | $F_{(1,10)}$ = 1.07 | P = 0.121 |
| Mid Hippocampus | AD | Genotype | $F_{(1,10)}$ <1 | P = 0.153 |
| | | Hemisphere | $F_{(1,10)}$ = 1.04 | P = 0.124 |
| Posterior Hippocampus | AD | Genotype | $F_{(1,10)}$ = 3.45 | P = 0.07 |
| | | **Hemisphere** | **$F_{(1,10)}$ = 6.19** | P = 0.052 |
| Anterior Hippocampus | RD | Genotype | $F_{(1,10)}$ = 3.03 | P = 0.086 |
| | | Hemisphere | $F_{(1,10)}$ <1 | P = 0.137 |
| Mid Hippocampus | RD | Genotype | $F_{(1,10)}$ <1 | P = 0.140 |
| | | Hemisphere | $F_{(1,10)}$ = 1.33 | P = 0.119 |
| Posterior Hippocampus | RD | **Genotype** | **$F_{(1,10)}$ = 10.83** | **P = 0.027** |
| | | Hemisphere | $F_{(1,10)}$ = 3.48 | P = 0.07 |

Statistical analysis of the anterior (Bregma -1.94 mm), mid (Bregma -2.46 mm) and posterior (Bregma -3.28 mm) hippocampus for fractional anisotropy (FA), apparent diffusion coefficient (ADC), axial diffusion (AD) and radial diffusion (RD). Analysis was performed using repeated measure two-way ANOVAs for genotype and hemisphere (Benjamini-Hochberg corrected (corrected P values stated)).

**Supp. Table 2**

| Brain Region | DTI Measure | ANOVA Comparison | F Value | P Value |
|---|---|---|---|---|
| Amygdala-Anterior Hippocampus | AD | Genotype | $F_{(1,10)}$ <1 | P = 0.164 |
| | | Hemisphere | $F_{(1,10)}$ = 2.10 | P = 0.097 |
| | | **Genotype x Hemisphere** | **$F_{(1,10)}$ = 12.12** | **P = 0.023** |
| Amygdala-Anterior Hippocampus | RD | Genotype | $F_{(1,10)}$ <1 | P = 0.149 |
| | | Hemisphere | $F_{(1,10)}$ = 1.32 | P = 0.106 |
| | | Genotype x Hemisphere | $F_{(1,10)}$ <1 | P = 0.155 |
| Amygdala-Posterior Hippocampus | AD | Genotype | $F_{(1,10)}$ <1 | P = 0.142 |
| | | Hemisphere | $F_{(1,10)}$ <1 | P = 0.189 |
| | | Genotype x Hemisphere | $F_{(1,10)}$ = 4.54 | P = 0.061 |
| Amygdala-Posterior Hippocampus | RD | Genotype | $F_{(1,10)}$ <1 | P = 0.151 |
| | | Hemisphere | $F_{(1,10)}$ <1 | P = 0.135 |
| | | Genotype x Hemisphere | $F_{(1,10)}$ <1 | P = 0.128 |
| BLA-Anterior Hippocampus | AD | Genotype | $F_{(1,10)}$ <1 | P = 0.167 |
| | | **Hemisphere** | **$F_{(1,10)}$ = 6.59** | **P = 0.047** |
| | | **Genotype x Hemisphere** | **$F_{(1,10)}$ = 10.53** | **P = 0.032** |
| BLA-Anterior Hippocampus | RD | Genotype | $F_{(1,10)}$ <1 | P = 0.158 |
| | | Hemisphere | $F_{(1,10)}$ = 2.59 | P = 0.092 |
| | | Genotype x Hemisphere | $F_{(1,10)}$ <1 | P = 0.173 |
| BLA-Posterior Hippocampus | AD | Genotype | $F_{(1,10)}$ <1 | P = 0.169 |
| | | **Hemisphere** | **$F_{(1,10)}$ = 12.79** | **P = 0.018** |
| | | **Genotype x Hemisphere** | **$F_{(1,10)}$ = 12.97** | **P = 0.02** |
| BLA-Posterior Hippocampus | RD | Genotype | $F_{(1,10)}$ <1 | P = 0.162 |
| | | Hemisphere | $F_{(1,10)}$ = 3.11 | P = 0.077 |
| | | Genotype x Hemisphere | $F_{(1,10)}$ <1 | P = 0.178 |

Statistical analysis of the anterior (Bregma -1.94 mm), and posterior (Bregma -3.28 mm) amygdala-hippocampal connections, analysed for axial diffusion (AD) and radial diffusion (RD). Analysis was performed using repeated measure two-way ANOVAs for genotype and hemisphere (Benjamini-Hochberg corrected (corrected P values stated)).

**Supp. Table 3**

| Brain Region | CLARITY Measure | ANOVA Comparison | F Value | P Value |
|---|---|---|---|---|
| M1 | OI | Genotype | $F_{(1,10)} <1$ | P = 0.182 |
| | | Hemisphere | $F_{(1,10)} = 1.74$ | P = 0.108 |
| M1 | Cell Density | Genotype | $F_{(1,10)} = 2.04$ | P = 0.099 |
| | | Hemisphere | $F_{(1,10)} = 1.41$ | P = 0.117 |
| M1 | Fibre Density | Genotype | $F_{(1,10)} <1$ | P = 0.171 |
| | | Hemisphere | $F_{(1,10)} <1$ | P = 0.176 |
| S1 | OI | Genotype | $F_{(1,10)} <1$ | P = 0.185 |
| | | **Hemisphere** | $\mathbf{F_{(1,10)} = 36.86}$ | **P = 0.005** |
| S1 | Cell Density | Genotype | $F_{(1,10)} <1$ | P = 0.131 |
| | | **Hemisphere** | $\mathbf{F_{(1,10)} = 13.73}$ | **P = 0.016** |
| S1 | Fibre Density | Genotype | $F_{(1,10)} = 1.73$ | P = 0.110 |
| | | **Hemisphere** | $\mathbf{F_{(1,10)} = 8.51}$ | **P = 0.038** |
| BF | OI | Genotype | $F_{(1,10)} <1$ | P = 0.191 |
| | | **Hemisphere** | $\mathbf{F_{(1,10)} = 10.59}$ | **P = 0.034** |
| BF | Cell Density | Genotype | $F_{(1,10)} <1$ | P = 0.133 |
| | | **Hemisphere** | $\mathbf{F_{(1,10)} = 8.70}$ | **P = 0.041** |
| BF | Fibre Density | Genotype | $F_{(1,10)} <1$ | P = 0.144 |
| | | Hemisphere | $F_{(1,10)} <1$ | P = 0.126 |

Statistical analysis of the primary motor cortex (M1), primary somatosensory cortex (S1) and the barrel field (BF). CLARITY imaged regions were then analysed for orientation index (OI), cell density and fibre density. Analysis was performed using repeated measure two-way ANOVAs for genotype and hemisphere (Benjamini-Hochberg corrected (corrected P values stated)).

**Supp. Figure 1**

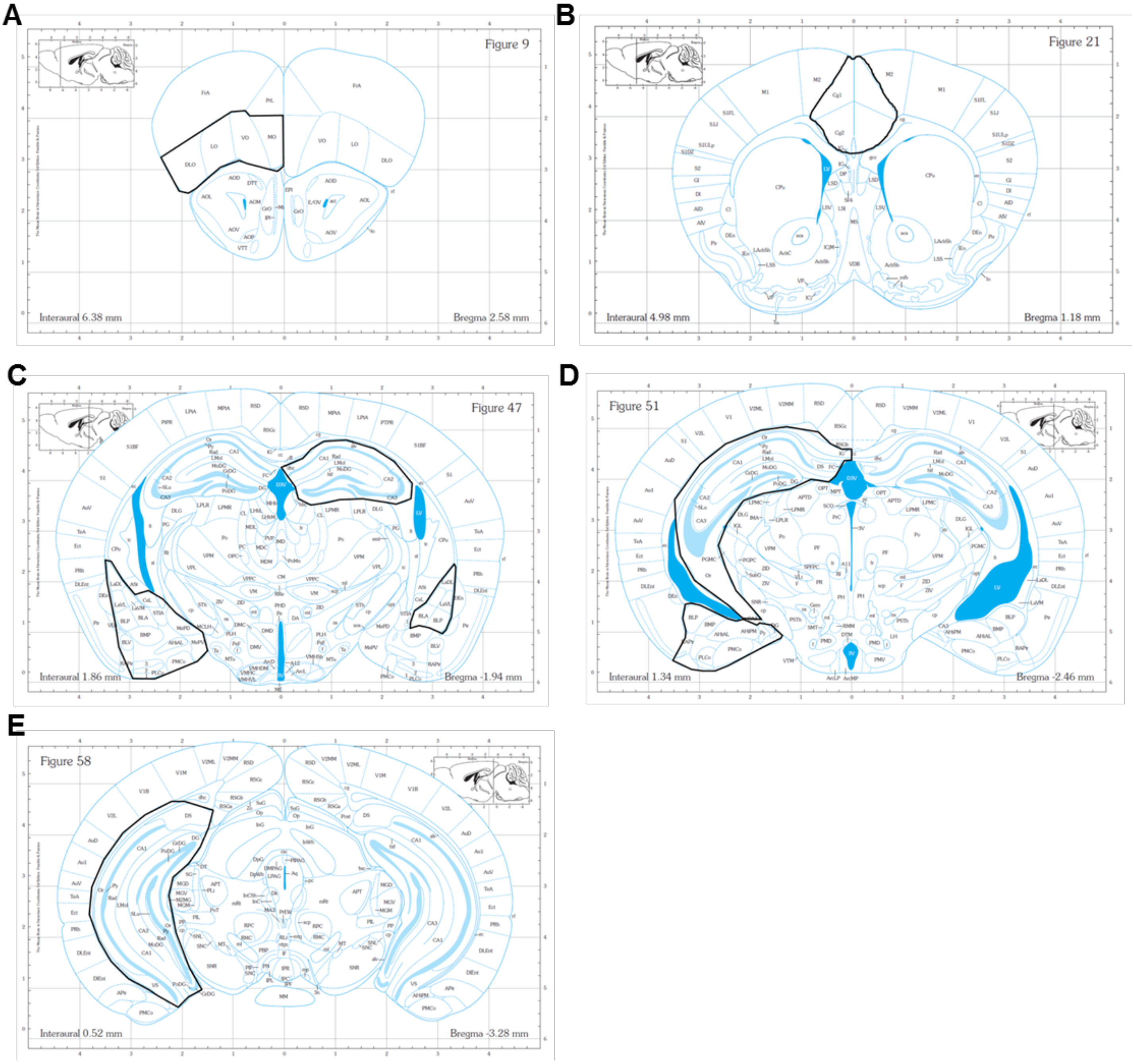


Atlas maps representing manual segmentation of regions of interest (ROI). (**A**) The orbitofrontal cortex ROI. (**B**) The ACC ROI. (**C**) The anterior hippocampus, anterior amygdala and basolateral amygdala ROI. (**D**) The mid hippocampus and posterior amygdala ROI. (**E**) The posterior hippocampus ROI. The atlas maps were used with the permission of the Authors (18).

**Supp. Figure 2**

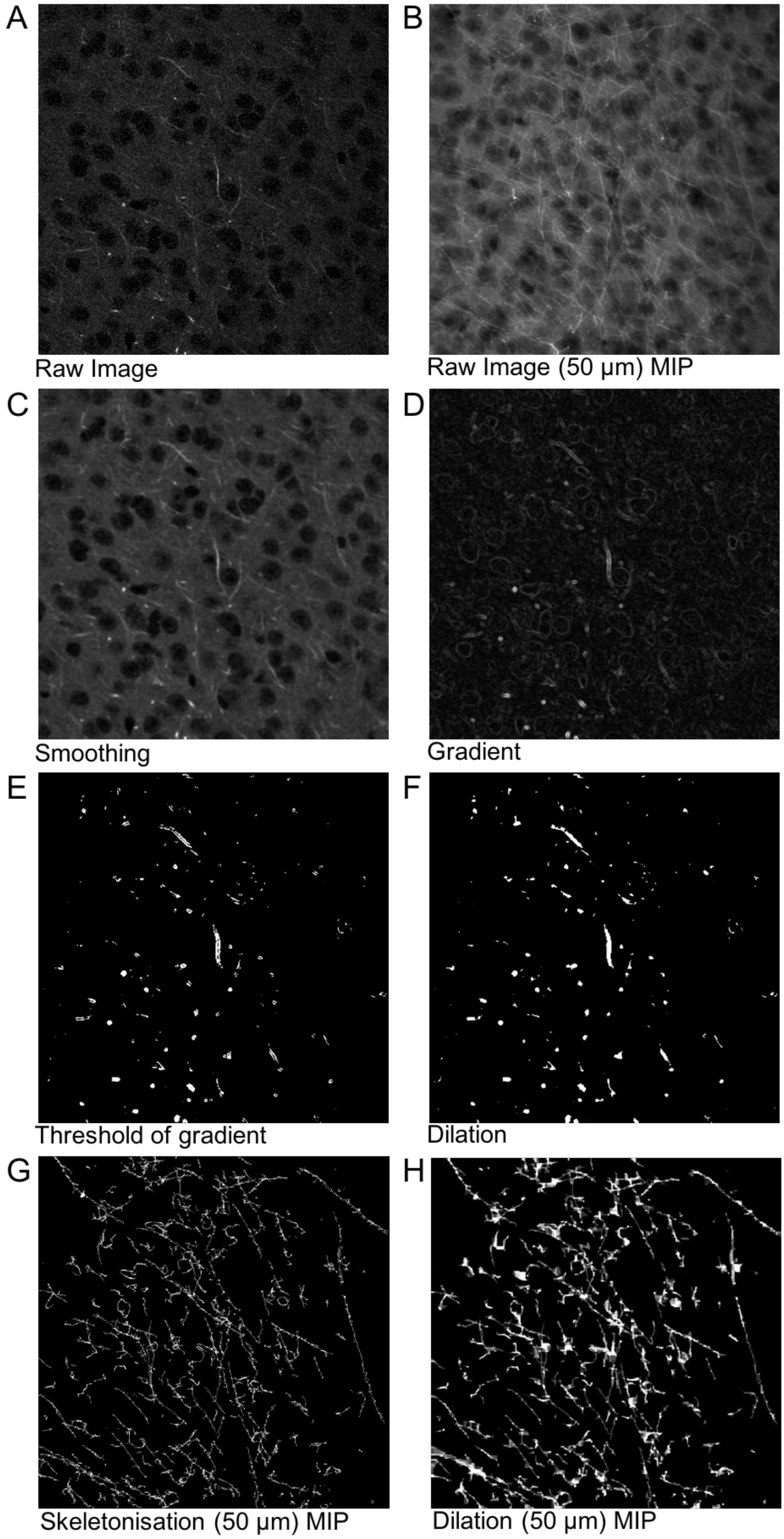


Analysis methodology of axonal segmentation from multiphoton images.

**Supp. Figure 3**

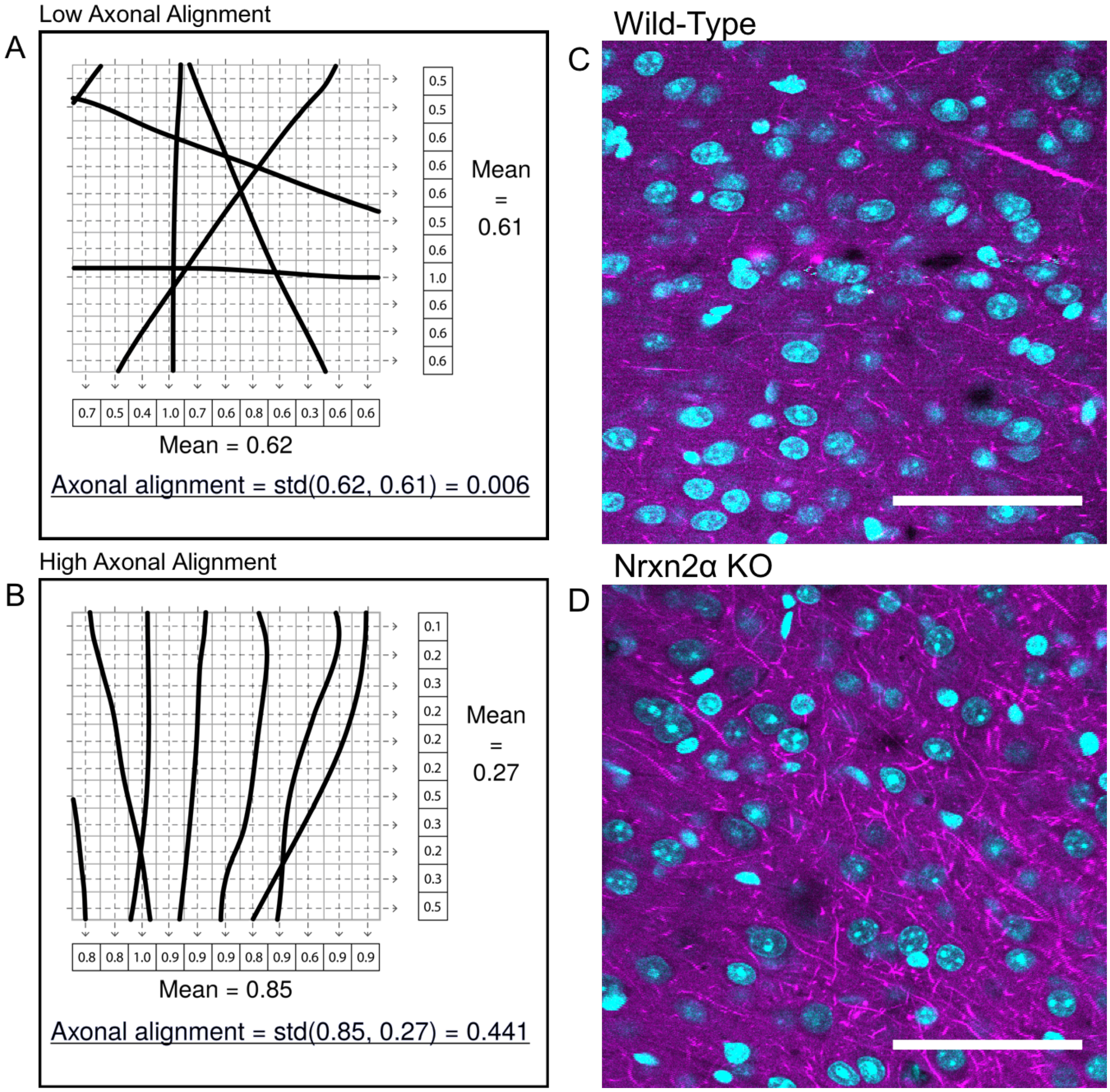


**A and B.** Illustration of the axonal alignment calculation in a simple two-dimensional case. The grey grid represents the image pixels, the black lines axons and the dashed lines represent the calculation process. Standard deviation denoted as std. **C and D.** Cingulate cortex images taken from wild-type and Nrxn2α KO mice, visually representing the greater axonal alignment and density in KO mice.

**Supp. Figure 4**

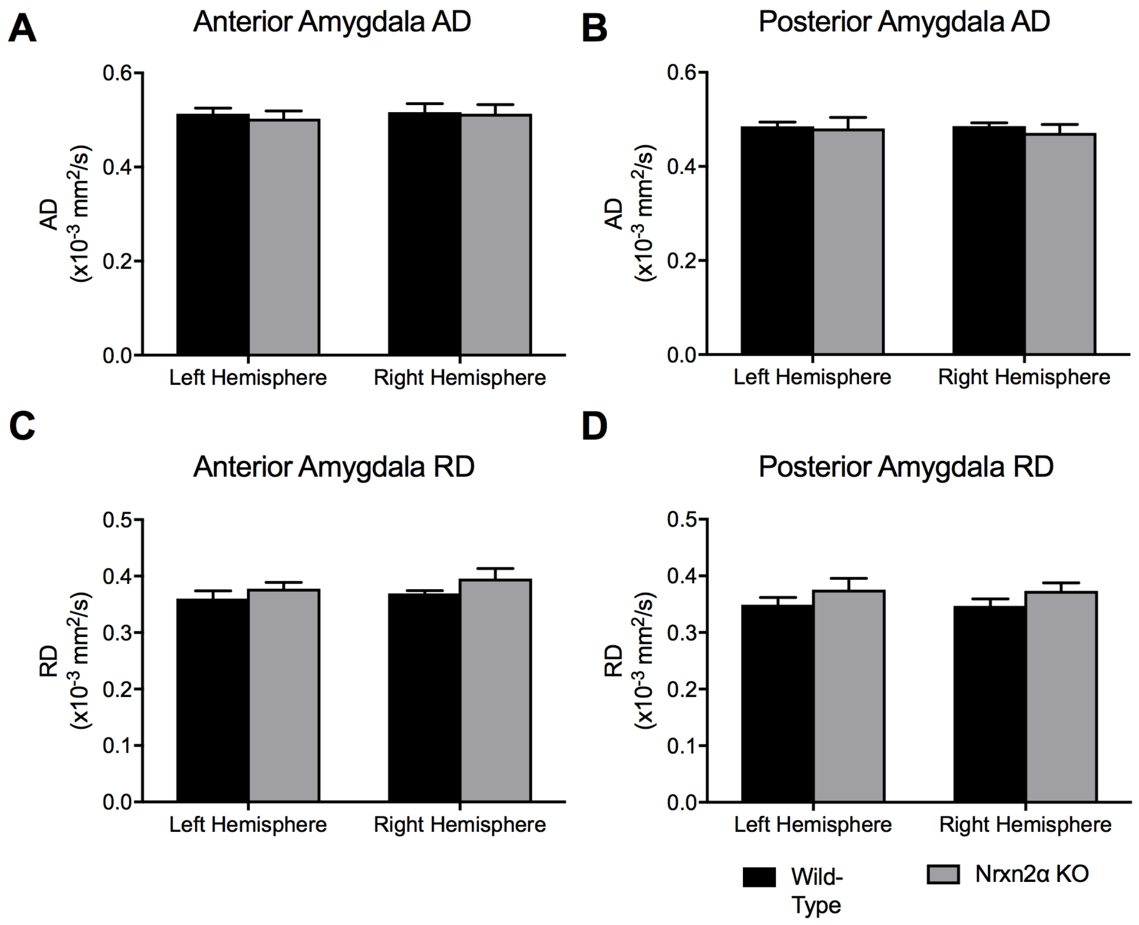


Axial diffusivity (AD) and radial diffusivity (RD) in the amygdala. The entire amygdala was segmented from DTI images at two regions; anterior (**A** and **C**: Bregma -1.94 mm) and posterior (**B** and **D**: Bregma -2.46 mm). No significant differences were observed between the genotypes for anterior or posterior regions. Error bars represent s.e.m. Wild-type n=6, Nrxn2α KO n=6.

**Supp. Figure 5**

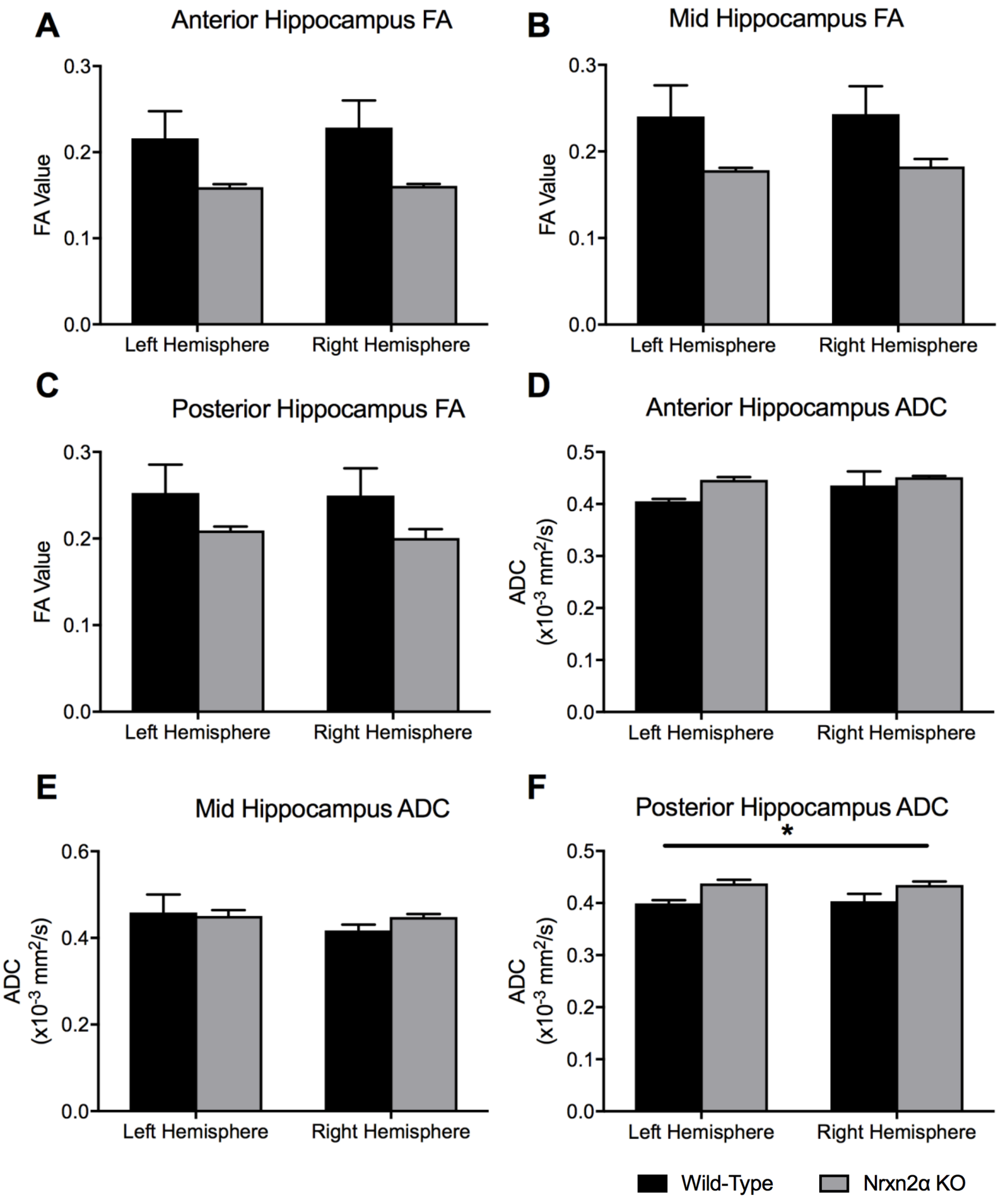


Fractional anisotropy (FA) and apparent diffusion coefficient (ADC) of the anterior to posterior hippocampus. Although Nrxn2α KO had trends towards lower FA in the anterior (**A**), mid (**B**) and posterior (**C**) hippocampus, there were no significant differences. Similarly, anterior (**D**) and mid (**E**) hippocampal regions did not vary between the genotypes, the posterior hippocampus (**F**) had significantly increased ADC (f). * = $P<0.05$. Error bars represent s.e.m. Wild-type n=6, Nrxn2α KO n=6.

**Supp. Figure 6**

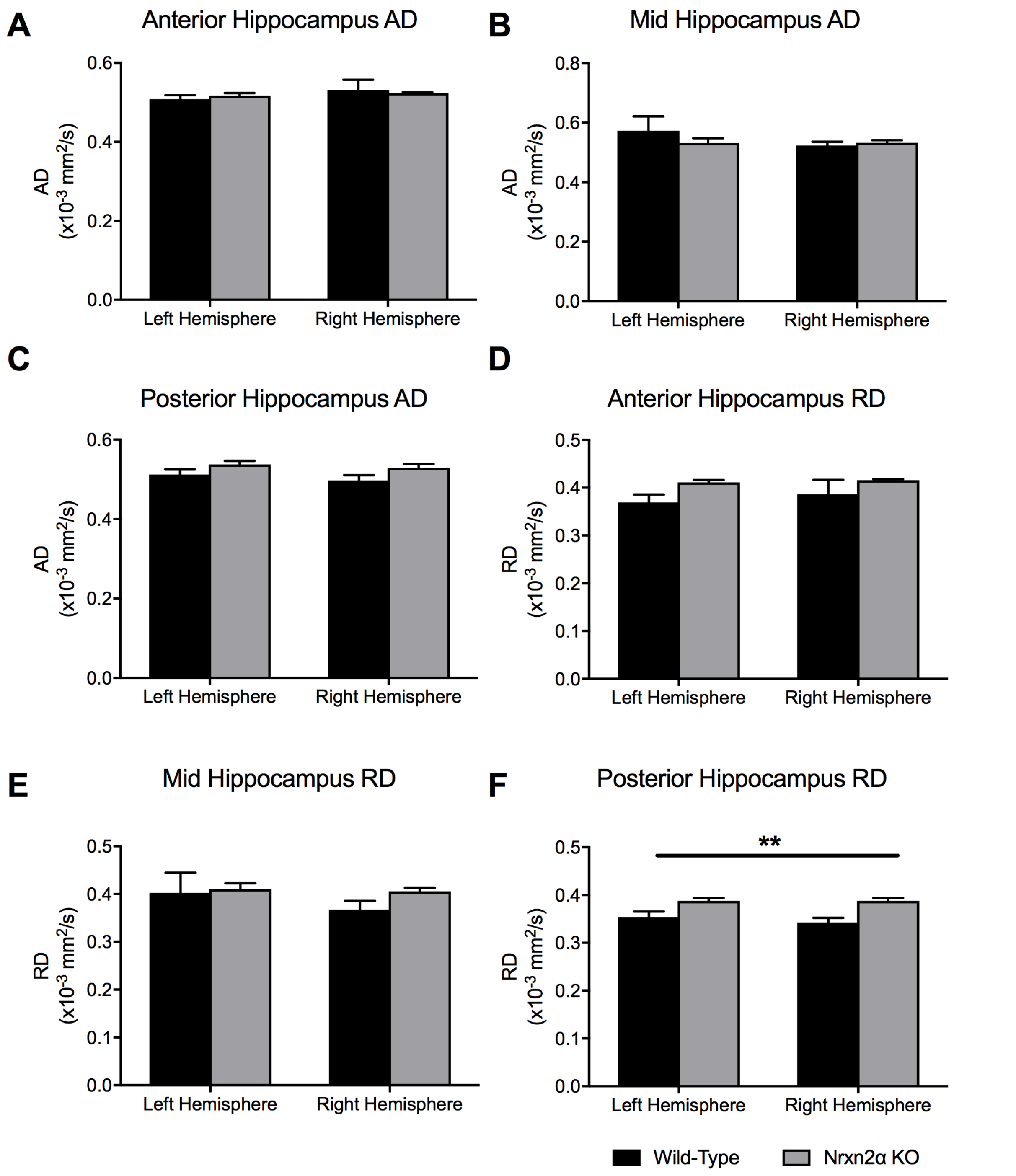


Axial diffusivity (AD) and radial diffusivity (RD) of the anterior to posterior hippocampus. AD did not differ in either the anterior (**A**), mid (**B**) and posterior (**C**) hippocampus. Although the anterior (**D**) and mid (**E**) hippocampal regions did not vary between the genotypes, the posterior hippocampus (**F**) had significantly increased RD (f). ** = $P<0.01$. Error bars represent s.e.m. Wild-type n=6, Nrxn2α KO n=6.

**Supp. Figure 7**

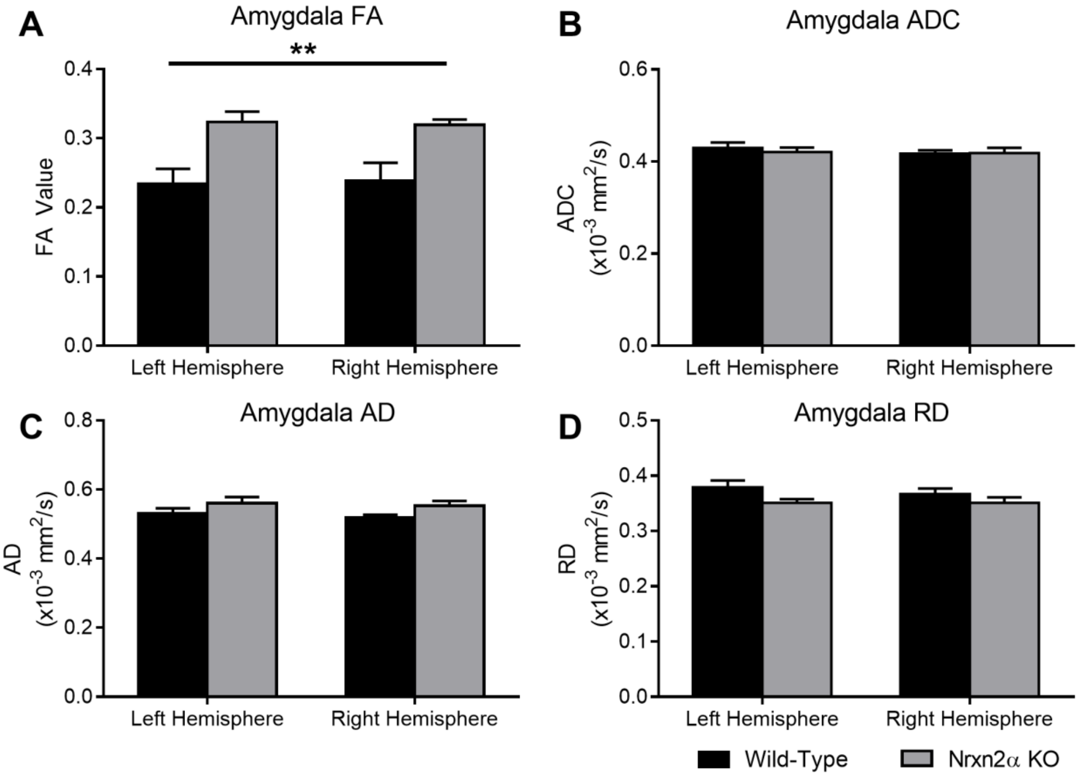


DTI quantified for the whole amygdala volume. (**A**) Fractional anisotropy (FA) was significantly increased in Nrxn2α KO mice (genotype: $F_{(1,10)} = 11.15$, $p = 0.008$). However, (**B**) apparent diffusion coefficient (ADC) was not significantly altered (genotype: $F_{(1,10)} <1$, $p = 0.782$), nor was (**C**) axial diffusivity (AD) (genotype: $F_{(1,10)} = 3.06$, $p = 0.111$) or (**D**) radial diffusivity (RD) (genotype: $F_{(1,10)} = 2.47$, $p = 0.147$). Error bars represent s.e.m. ** = P<0.01. Wild-type n=6, Nrxn2α KO n=6.

**Supp. Figure 8**

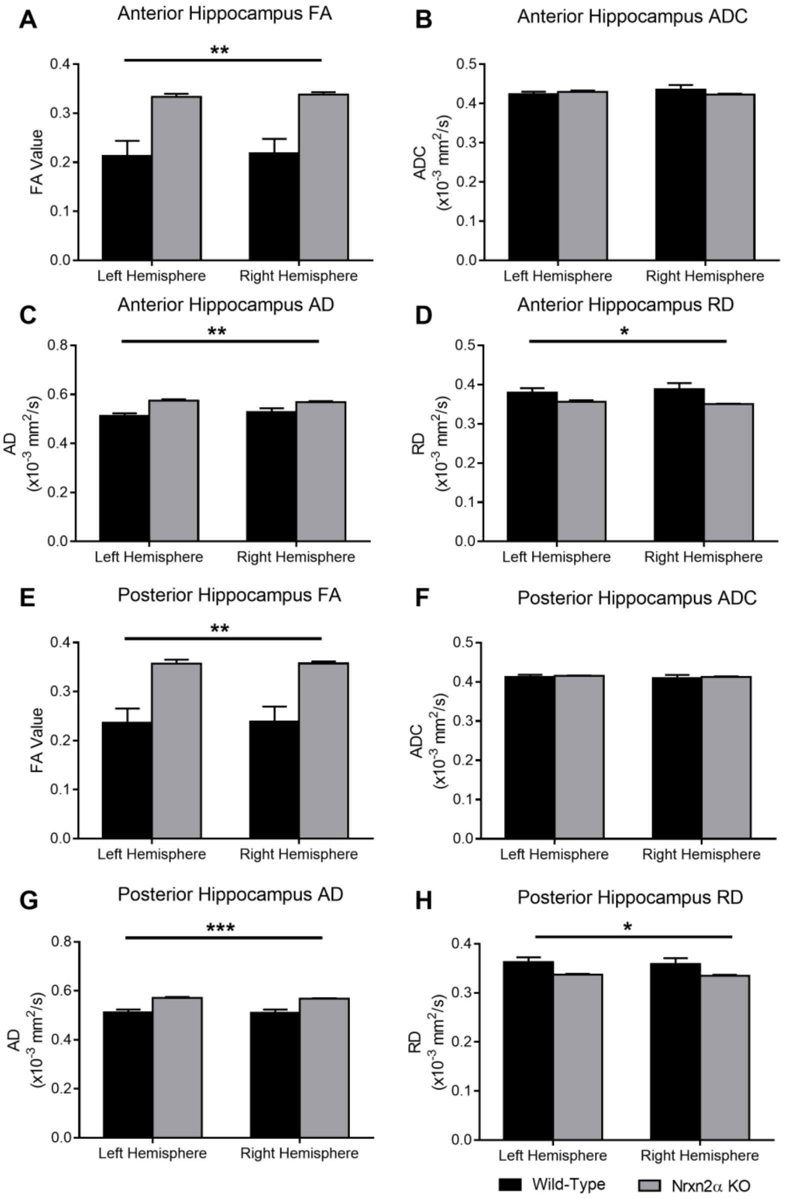


DTI quantified for the whole anterior hippocampus (Bregma -1.06 mm – -2.46 mm) and posterior hippocampus (Bregma -2.54 mm – -3.16 mm). (**A**) Fractional anisotropy (FA) in the anterior hippocampus was significantly increased in Nrxn2α KO mice (genotype: $F_{(1,10)}$ = 15.63, p = 0.0027) but (**B**) apparent diffusion coefficient (ADC) was not altered (genotype: $F_{(1,10)}$ <1, p = 0.738). (**C**) Axial diffusivity (AD) (genotype: $F_{(1,10)}$ = 16.17, p = 0.0024) and (**D**) radial diffusivity (RD) (genotype: $F_{(1,10)}$ = 5.05, p = 0.048) were both significantly altered in Nrxn2α KO mice. In the posterior hippocampus, in Nrxn2α KO mice, (**E**) FA was significant increased (genotype: $F_{(1,10)}$ = 15.62, p = 0.0027), (**F**) ADC was similar to wild-types (genotype: $F_{(1,10)}$ <1, p = 0.679), (**G**) AD was increased (genotype: $F_{(1,10)}$ = 22.31, p = 0.0008) and (**H**) RD was significantly reduced (genotype: $F_{(1,10)}$ = 5.34, p = 0.043). Error bars represent s.e.m. * = P<0.05, ** = P<0.01, *** = P<0.001. Wild-type n=6, Nrxn2α KO n=6.

**Supp. Figure 9**

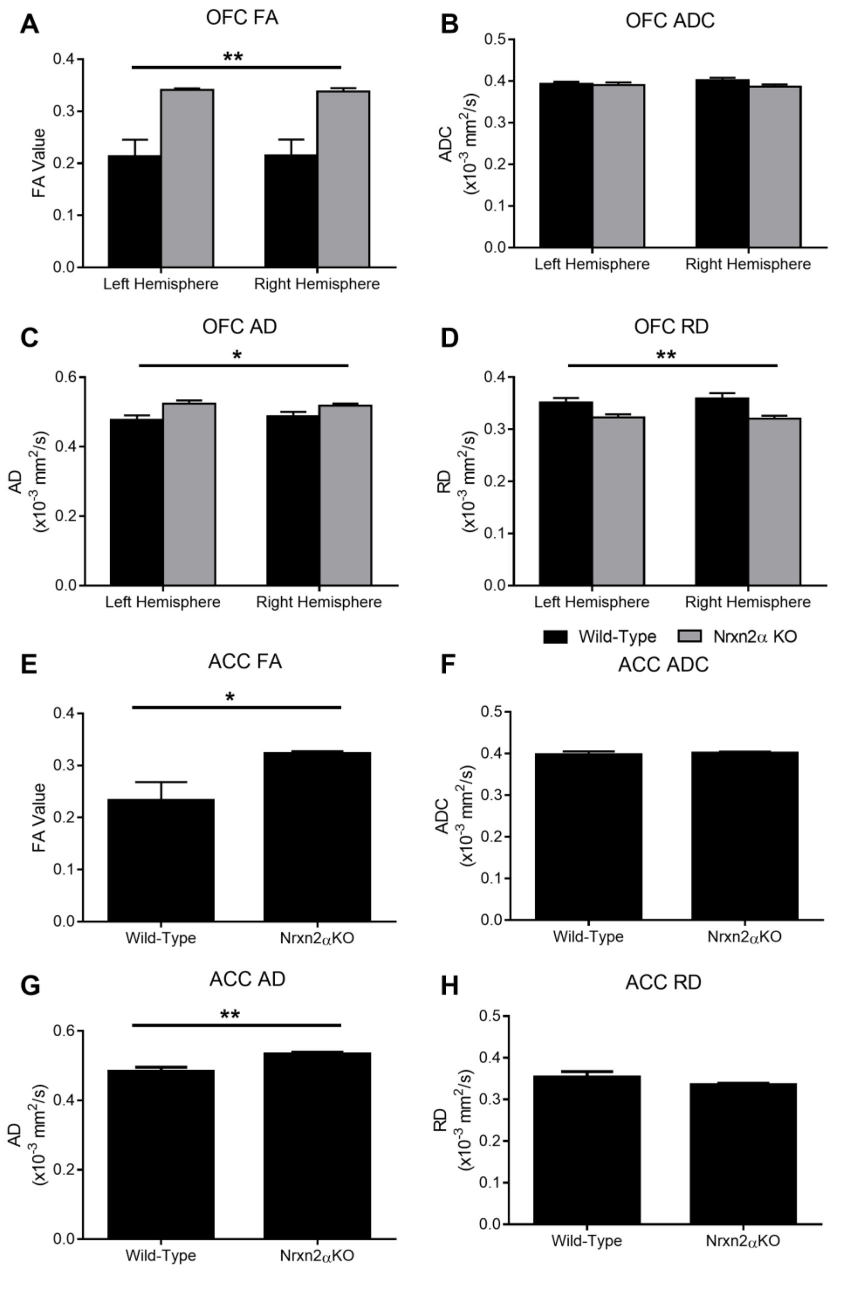


DTI quantified for the whole orbitofrontal cortex (OFC) and anterior cingulate cortex (ACC) and posterior hippocampus (Bregma -1.34 mm – -0.46 mm). (**A**) Fractional anisotropy (FA) in the OFC was significantly increased in Nrxn2α KO mice (genotype: $F_{(1,10)}$ = 16.14, p = 0.0024) but (**B**) apparent diffusion coefficient (ADC) was not altered (genotype: $F_{(1,10)}$ = 1.43, p = 0.260). (**C**) Axial diffusivity (AD) (genotype: $F_{(1,10)}$ = 6.71, p = 0.027) and (**D**) radial diffusivity (RD) (genotype: $F_{(1,10)}$ = 10.07, p = 0.0099) were both significantly altered in Nrxn2α KO mice. In the ACC of Nrxn2α KO mice, (**E**) FA was significant increased (genotype: $t_{(10)}$ = 2.55, p = 0.029), (**F**) ADC was similar to wild-types (genotype: $t_{(10)}$ = 2.55, p = 0.618), (**G**) AD was increased (genotype: $t_{(10)}$ = 3.89, p = 0.003) and (**H**) RD was not significantly different (genotype: $t_{(10)}$ = 1.35, p = 0.208). Error bars represent s.e.m. * = P<0.05, ** = P<0.01. Wild-type n=6, Nrxn2α KO n=6.

**Supp. Figure 10**

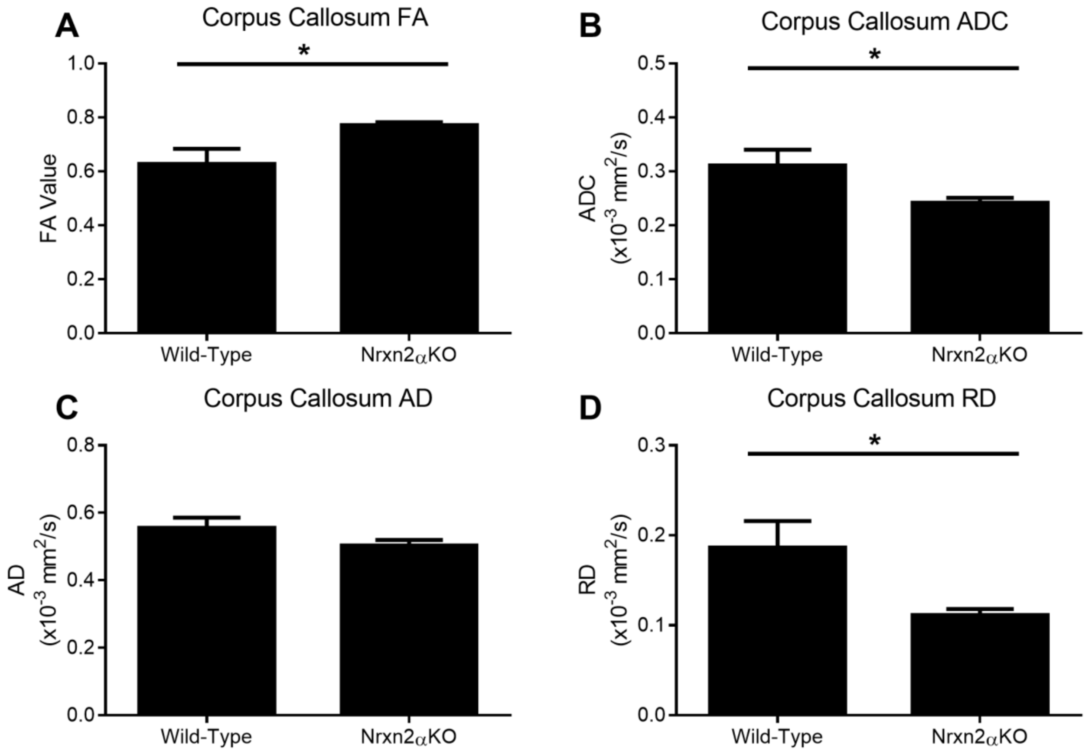


DTI quantification of the corpus callosum (Bregma 0.98 mm). To examine the integrity of white matter tracts within Nrxn2α KO mice, we examined diffusivity in the corpus callosum. (**A**) Fractional anisotropy (FA) was significantly increased in Nrxn2α KO mice (genotype: $t_{(10)} = 2.50$, $p = 0.032$) and apparent diffusion coefficient (ADC) (**B**) significantly decreased (genotype: $t_{(10)} = 2.28$, $p = 0.046$). This difference appeared to be driven predominantly by radial diffusivity (RD), as axial diffusion (AD) (**C**) was not significantly different (genotype: $t_{(10)} = 1.49$, $p = 0.168$) whilst RD (**D**) was significantly reduced in Nrxn2α KO mice (genotype: $t_{(10)} = 2.45$, $p = 0.034$). Error bars represent s.e.m. * = $P<0.05$. Wild-type n=6, Nrxn2α KO n=6.

**Supp. Figure 11**

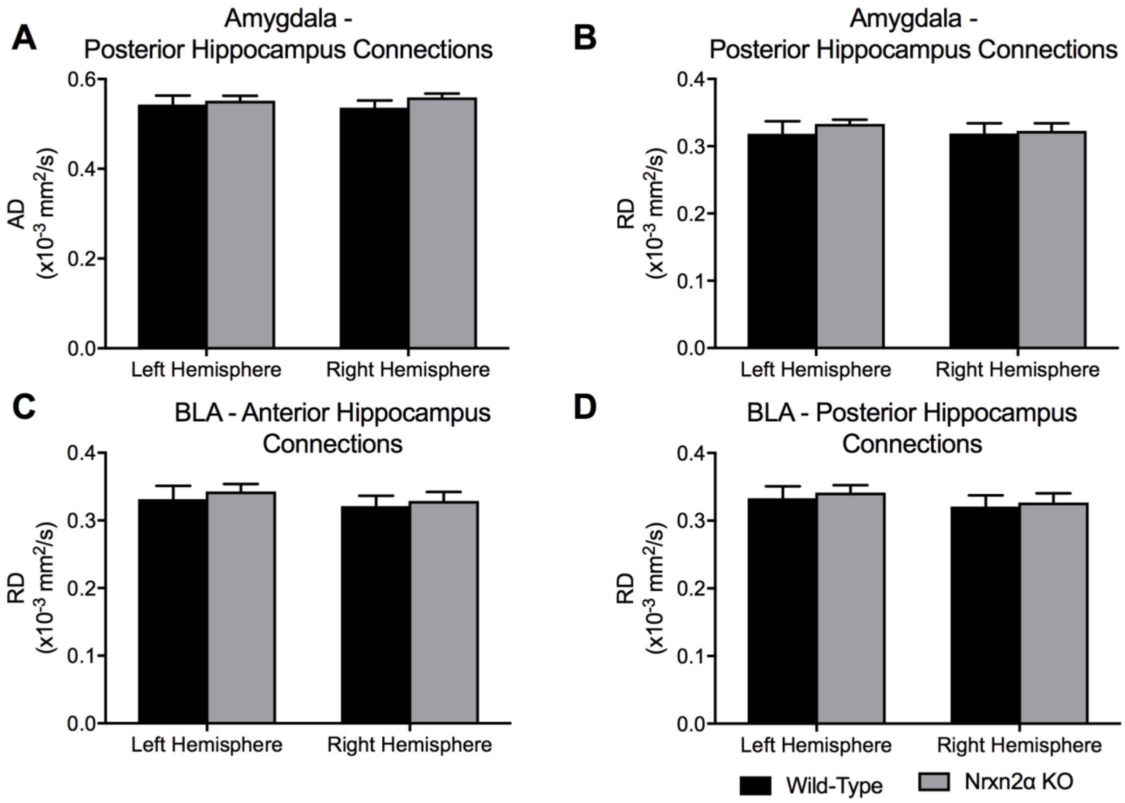


Axial diffusivity (AD) and radial diffusivity (RD) of computed tracts of connections from the amygdala to hippocampus. Tracts from the anterior amygdala to the posterior hippocampus (Bregma -2.46 mm) were analysed for AD (**A**) and RD (**B**). No significant differences between the tracts of Nrxn2α KO mice were observed. No significant differences were found for RD of tracts specifically from the basolateral nuclei of the amygdala (BLA) to the anterior (**C**) or posterior (**D**) hippocampus. Error bars represent s.e.m. Wild-type n=6, Nrxn2α KO n=6.

**Supp. Figure 12**

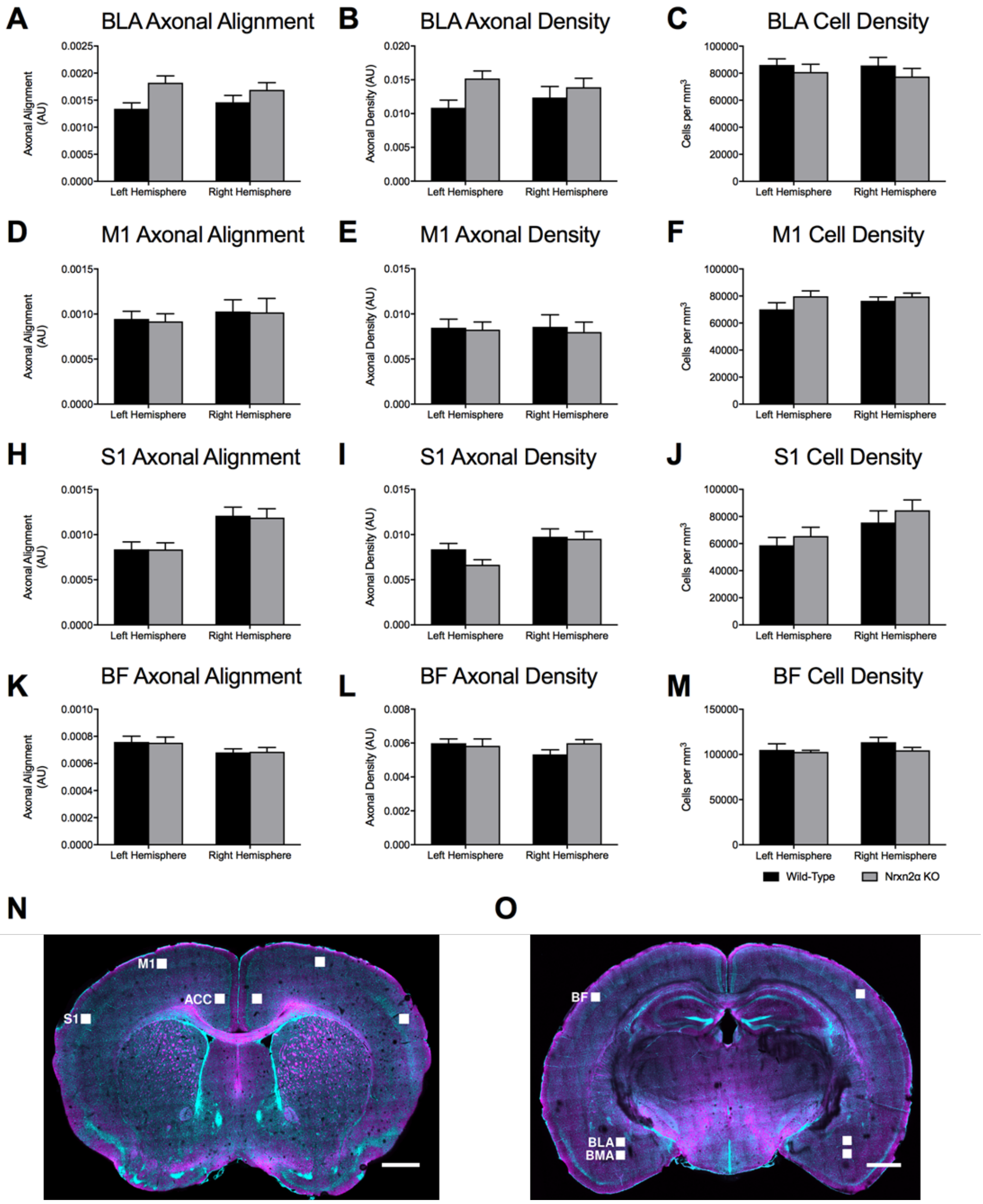


CLARITY-derived quantification of fibres and cell density within the basolateral amygdala (BLA) and control regions (**A**) Although there were trends towards increased axonal alignment and fibre density (**B**) in Nrxn2α KO mice, no significant differences were found. (**C**) Cell density in the BLA was similar between the genotypes. Statistical analysis (Supp. Table 4) was performed for the primary motor cortex (M1; **D-F**), primary somatosensory cortex (S1; **H-J**) and the barrel field (BF; **K-M**). No genotypic differences were found for any measure within these cortical regions. (N-O) CLARITY images of the scanned regions of interest. Error bars represent s.e.m. Wild-type n=6, Nrxn2α KO n=6.
